# Human social learning biases in immersive virtual environments

**DOI:** 10.1101/2022.08.26.504784

**Authors:** C. Easter, C. Hassall

## Abstract

Social learning strategies describe what, when, and from whom individuals choose to learn. Evidence suggests that both humans and animals are capable of strategic social learning. However, human research generally lacks ecological and spatial realism, making it difficult to understand the importance of our use of social information in an evolutionary context. In this study, we use virtual reality to simulate three novel tasks inspired by the animal literature (Container, Route Choice and Foraging tasks) within complex, three-dimensional environments. In each experiment, combinations of demonstrators with different characteristics gave opposing solutions to the task to determine from whom participants preferentially learned. Importantly, participants were able to freely navigate the environment and attempt the task in any way they chose by using or ignoring social information. We found that participants displayed an overall bias towards learning asocially (independently) rather than socially. Asocial learning was favoured more strongly during complex tasks that spanned larger spatial scales, potentially due to the difficulties in keeping track of social information in such scenarios. When learning from others, participants displayed a bias towards learning from the majority over the minority (positive frequency-dependent social learning) and towards learning from the most successful demonstrators (payoff-based social learning), which supports the findings of previous, lab-based experiments. There was no apparent bias with respect to demonstrator dominance status, gender and body size. Our findings are the first to show a variation in the use of social learning across task and environmental complexities in humans, to carry out a comprehensive evaluation of hypothetical human learning biases, and to provide a methodological link between non-human and human social learning experiments. As demonstrated here, immersive virtual environments have great potential for research into human social evolution and we strongly encourage future research to adopt a similar approach.

## Introduction

‘Social learning strategies’ describe how individuals choose to learn from one another – specifically when they favour social over asocial (independent) learning, what type of information or behaviour they learn, and from whom they learn (Laland, 2004; Rendell *et al*., 2011). Empirical research suggests that both humans and non-human animals are strategic in their use of social information – e.g. by copying when asocial learning is costly (Morgan *et al*., 2012; Webster and Laland, 2008; Coolen, *et al*. 2003), copying when the task is more difficult (Flynn *et al*., 2016) or selectively copying the most successful and/or proficient individuals (Mesoudi, 2008; Mesoudi and O’Brien, 2008; Morgan *et al*., 2012; Kendal, Rendell, *et al*., 2009; Pike *et al*., 2010; Coelho *et al*. 2015). Both humans and non-human animals also display ‘model-based’ learning biases towards individuals with certain characteristics. For example, various species, including humans and certain primates, display biases towards copying high-ranking and/or older individuals (Henrich and Henrich, 2010; Horner *et al*., 2010; Coelho *et al*. 2015; Kendal *et al*., 2015), potentially because these characteristics act as an indicator of an individual’s overall success. There is also evidence across taxa that individuals tend to conform to the majority or group norms (Efferson *et al*., 2008; Morgan *et al*., 2012; Pike and Laland, 2010; Hopper *et al*. 2011; Van de Waal *et al*., 2013). For humans, in particular, social learning has been paramount to our development of culture.

Considering human social learning strategies from an evolutionary perspective is, however, difficult due to the highly abstract laboratory experiments often used in human research. Where animal research generally involves investigating how freely interacting individuals learn from one another while completing ecologically relevant tasks such as finding food or avoiding predators, human experiments tend to lack realism for three main reasons. Firstly, human experiments often involve the solving of highly abstract tasks with little ecological significance (e.g. choosing between coloured options, Efferson *et al*., 2008; mentally rotating objects, Morgan *et al*., 2012; or building model towers, Caldwell and Eve, 2014). Secondly, social information tends to be represented in an overly simplistic way – e.g. as numerical values (Mesoudi and O’Brien, 2008) or flashing tiles (Morgan *et al*., 2012) – while, in reality, individuals would need to actively observe the behaviours of others in order to learn from them. Thirdly, human experiments are almost always restricted to the laboratory, and so confined to extremely small spatial scales – despite the fact that many human social learning processes, now and in our evolutionary past, generally occur over very large spatial scales (Hamilton *et al*., 2007; Whallon, 2006). For these reasons, it can be difficult to consider human social learning strategies in a common evolutionary framework with animal behavioural research.

Virtual reality (VR) offers one largely unexploited way to study human social behaviour in more ‘realistic’ settings. In particular, large, open-world virtual environments in which individuals can freely explore, learn and interact while completing complex, ecologically relevant tasks across large spatial scales could act as a unique testbed for human social learning research. Currently, massively multiplayer online roleplaying games (MMORPGs) such as *World of Warcraft* offer the most potential for studying human social dynamics within virtual spaces (Lofgren and Fefferman, 2007). However, commercial games like these have not been purpose built for behavioural research. Here, we introduce a novel research tool specifically designed for the purpose of conducting social learning experiments within realistic virtual spaces and conduct experiments inspired by the animal literature to test key hypotheses concerning human social information use within ecologically and spatially realistic scenarios. We focus on (a) whether participants preferentially learn socially or asocially when exposed to a series of novel tasks of varying complexity; (b) whether participants display biases towards learning from demonstrators with particular physical and/or behavioural characteristics, and (c) whether participants are more prone to copying demonstrators that use more rewarding behaviours. We discuss these findings in relation to previous work on humans conducted in more restrictive laboratory settings and also in relation to the animal literature. By generating an experimental environment where participants have the freedom to navigate large-scale landscapes and complete tasks as they wish, and where the use of social information requires the active observation of knowledged demonstrators, this study also aims to bridge the gap between animal and human research by demonstrating how virtual reality can offer a highly promising way of studying human social behaviour within an evolutionary framework.

## Methods

To evaluate the role that demonstrator characteristics play in mediating the use of social information, we developed a new virtual laboratory: Virtual Environments for Research into Social Evolution (VERSE). VERSE allows researchers to create realistic, large-scale three-dimensional environments containing ecologically relevant tasks that allow direct comparison to (or replication of) behavioural experiments on freely interacting animal populations. Within VERSE, participants take control of a virtual ‘player’ to explore and interact with the environment freely. Artificial intelligence agents (AIs) can be added to the environment and programmed with specific behaviours such as random walks, route following and object interactions to provide sources of social information. Crucially, if participants are to take advantage of this social information, they must observe AI behaviours and decide which aspects of those behaviours to copy – thus, VERSE immerses participants in a realistic social environment and avoids the oversimplification of social information that is often present in lab studies. For more details about VERSE and its features, see Easter (2022).

### General experimental design

One hundred and forty-two participants (51 males, 91 females; ages 18-31 years; mean age = 20.6 years) controlled a virtual human ‘player’ within a number of realistic, three-dimensional virtual environments (constructed using VERSE) in a series of three experiments inspired by the animal literature. All participants were undergraduates from the Faculty of Biological Sciences, University of Leeds and were recruited via email advertisements. Prior to starting the study, participants were given instructions on how to control the virtual player, including how to move the player, rotate the camera and interact with objects, and were then asked to practice this within a ‘demo’ environment. This practice phase ensured that all participants were comfortable with the game controls prior to starting the experiment, regardless of prior IT and/or gaming experience. All participants received a £15 Amazon voucher as compensation for their time. Ethical approval for this study was obtained by the Faculty of Biological Sciences Research Ethics Committee (LTSBIO-029).

Each experiment required participants to complete a novel task (detailed below) in the presence of computer-controlled AI ‘demonstrators’, which acted as realistic sources of social information. Each task was replicated across six ‘demonstrator conditions’, designed to test key hypotheses about social learning biases highlighted throughout the literature. For each demonstrator condition, two sets of humanoid AI ‘demonstrators’ with different physical and/or behavioural characteristics (hereon referred to as ‘demonstrator A’ and ‘demonstrator B’) were programmed to display different solutions to the given task. Participants were notified of the presence of ‘other players’ who they were not in competition with and who had prior knowledge of the task. They were not informed whether they were expected to use information supplied by these players. No restrictions were placed on when and from whom social information could be used – participants were free to watch, copy or ignore demonstrators entirely. The order of presentation for both tasks and demonstrator conditions was randomised across participants according to a balanced Latin square design (Supplementary Material, Tables S1-S3).

The demonstrator conditions were as follows. ***Asocial***: No demonstrators present. ***Soc/Asoc*:** Demonstrator A is a single AI; no alternative demonstrator is present. ***Dom/Sub*:** Demonstrator A is one ‘dominant’ AI with a muscular appearance; demonstrator B is one ‘subordinate’ AI with a hunched, emaciated appearance. At the beginning of the task, the dominant AI additionally displays an aggressive display (punching outwards) towards the subordinate AI, and the subordinate responds by shielding its face and turning away. ***Three/One*:** Demonstrator A is three AIs all performing the same solution at the same time; Demonstrator B is a single AI. ***Male/Female*:** Demonstrator A is one ‘male’ AI (with default AI appearance); demonstrator B is one ‘female’ AI (modified to enhance the waist-to-hip ratio and reduce muscle mass). ***Large/Small*:** Demonstrator A is an AI larger than the player; demonstrator B is an AI smaller than the player. Prior to the main experiment, 41 participants completed a questionnaire regarding their perceptions of the AI models to be used in the main study to explore whether the models to be used were perceived in the intended way (e.g. the ‘female’ AI as female, the ‘dominant’ AI as dominant to the ‘subordinate’ AI, etc.). Results from this validation exercise are discussed in more detail in the Supplementary Material (Tables S4-S5) but suggest that people perceived the AIs in the intended way. Participants were assigned to one of two ‘reward groups’ designed to investigate how demonstrator success (in addition to their individual characteristics) influenced their likelihood of being copied. In the ‘Same Rewards’ group (*n* = 72), the choices made by both demonstrators in each demonstrator condition were equally profitable and so participants received the same in-game payoff regardless of which demonstrator they copied. This condition was used to investigate participants’ innate biases towards particular demonstrators when there was no direct benefit of choosing to learn from specific individuals. In the ‘Different Rewards’ group (*n* = 71), different solutions to a task resulted in a different payoff. For each demonstrator condition, demonstrator A always provided a higher payoff solution to each task than demonstrator B, thus giving participants the opportunity to learn which demonstrators tended to be the most successful. Comparisons of the two reward groups were made to investigate whether participants tended to learn from more successful demonstrators, beyond any innate preferences they may hold for demonstrators with particular characteristics.

Game data was collected automatically during gameplay. After the completion of all three tasks, participants uploaded their game data anonymously to a file request link and filled in a post-experiment questionnaire (detailed below).

### Tasks

#### Container task

This task (inspired by Horner *et al*.’s 2010 experiment on chimpanzees) was designed to test the influence of different demonstrators on participant choices when making simple, binary decisions. Participants were required to pick up and deposit a token into one of two possible containers – blue(/left) or yellow(/right) – over 10 rounds per demonstrator condition to receive food rewards (Figure 1A-C). Participants were instructed to gather as many food rewards as they could. They were informed that the rewards varied between rounds and that the number of rewards they received would depend on their choice of container. This task was replicated once per demonstrator condition (i.e. six game levels in total) after which the participant was then given their ‘food collection score’ (percentage of possible food items collected) before moving to the next level / demonstrator condition. For the Same Rewards group, the participant received the same number of food items regardless of which container they chose on a particular round. For the Different Rewards group, the number of food items dispensed differed between containers. At the start of each round, one container was randomly selected as the ‘best’ choice and was allocated a random reward number between four and six. The other container was selected as the ‘worst’ choice and was allocated a random reward number between one and three. For each demonstrator condition, demonstrator A always chose the container with the best reward.

**Figure 1:**
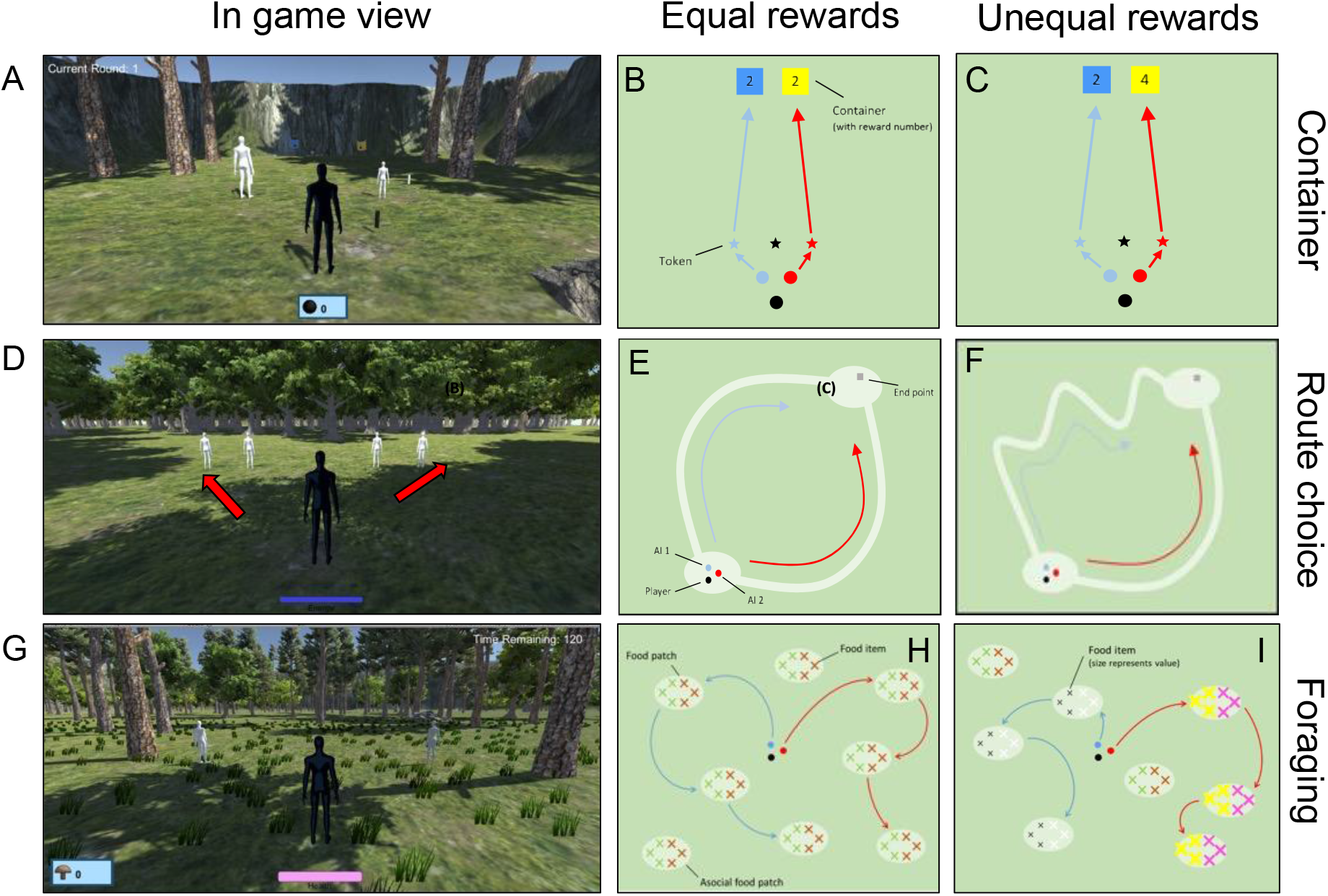
Three virtual reality tasks showing in-game views (A,D,G), top-down schematics of the game space with illustration of equal rewards (B,E,H), and top-down schematics showing unequal rewards depending on choices (C,F,I). See text for details.

##### Data collection and preparation

The container chosen by the player and each demonstrator during each round were logged. A participant was said to have copied a particular demonstrator if they chose the same container as that demonstrator during a given round. Since there were only two possible choices in this task, participants were therefore forced to choose the same container as one of the demonstrators (with the exception of the *Soc/Asoc* condition, where one option was always undemonstrated). As such, there was no scope for asocial learning that was completely independent of demonstrator choices. For each participant, the total number of times each demonstrator was copied during each demonstrator condition was then calculated.

#### Route Choice task

This task (inspired by Laland and Williams’ 1998 experiment on guppies) was designed to test how human route choice preferences within naturalistic, large-scale environments were influenced by the choices of other individuals. Participants were placed in a series of forest environments and instructed to find their way through the forest to a hidden cave using as little ‘energy’ as possible. In each environment, two clear, treeless paths were available (left or right), both of which led to the endpoint (Figure 1D-F). Three replicates were used for each of the six demonstrator conditions (18 game levels in total). Each replicate consisted of a new, visually distinct environment, with the endpoint and two clear paths positioned differently. Participants began each level with an energy value of 100% and the player could see their energy depleting during movement. For the Same Rewards group, the length of each clear path, and so the energy required to navigate them, was approximately the same (Figure 1E). One demonstrator was randomly chosen for each path. For the Different Rewards group, one path was substantially longer, and so more energetically costly to travel, than the other (Figure 1E), and demonstrator A always chose the shorter path. The direction (left or right) of the shortest path was randomised between replicates to prevent participants basing their choices purely on direction. Participants were given complete freedom of movement – they could choose to follow the demonstrators, choose one of the paths independently, or ignore the clear paths entirely and attempt to navigate the trees instead. Thus, participants had the opportunity to use social information or make completely independent decisions.

##### Data collection and preparation

The locations of the player and all AIs were logged at 1s intervals to map the exact routes taken by each character. A 15-unit buffer, which approximately matched the width of the clear paths, was created around each demonstrator’s route using the gBuffer() R function (*rgeos* package; Bivand and Rundel, 2020). Using every fifth data point from the player’s location data, the point.in.polygon() R function (*sp* package; Pebesma and Bivand, 2005) was used to calculate how many of the player’s location points fell within the buffer region of each demonstrator’s route. If ≥50% of the player’s route was contained within the buffer region of a particular demonstrator’s route, the participant was considered to have copied the route choice of that demonstrator. Otherwise, the participant was considered to have navigated the forest independently. For each participant, the total number of times each demonstrator was copied and the total number of times a route was chosen asocially were calculated.

#### Foraging task

This task (inspired generally by experiments on free-roaming animal populations) was designed to test the influence of different demonstrators on participant behaviour in a realistic foraging scenario. Participants were given 200 seconds to explore a large, open environment in search of food (Figure 1G-I). There was one replicate per demonstrator condition (i.e. six game levels in total), each with a different environment. Within each environment, there were 8 randomly located food patches, each containing a total of 40 food items (two sets of 20, where each set was a different ‘food type’). Food items all looked like mushrooms, but different food types were coloured differently and, in some cases, had different nutritional values (Supplementary Material, Table S6). The player was given a food score and health bar, which were displayed onscreen. When the player collected a food item, its nutritional value was added to their food score. Food could also be poisonous (i.e. have a negative nutritional value). Collecting poisonous food reduced the player’s food score and health. If the player’s health reached zero due to eating poisonous food items, the current level ended prematurely. Participants were informed of this prior to starting the task. Once 200 seconds had elapsed, participants were given their food score as a percentage of the maximum score that could have been obtained if they had collected all non-poisonous foods in the area, before proceeding to the next level. For each demonstrator condition, three of the eight available food patches were visited by demonstrator A, another three were visited by demonstrator B and the remaining two were not visited by any demonstrator (i.e. they were ‘asocial patches’ that could only be discovered independently). Each demonstrator visited their chosen food patches in a random order. Once they reached a food patch, demonstrators would ‘eat’ 10 food items from only one of the available food types located there, thus demonstrating food preferences to the participant while leaving the majority of food items for the player to collect. For the Same Rewards group, all food patches contained the same two food types, each of which had the same nutritional value of +1 (Figure 1H). For the Different Rewards group, each food patch contained different food types with different nutritional values (Figure 1I). The three food patches visited by demonstrator A contained two food types with the highest nutritional values, +5 and +3. The three food patches visited by demonstrator B contained two food types with the lowest nutritional values, +1 and -1. The latter was a poisonous food type which depleted the player’s health and food score. In each case, the demonstrator only ‘ate’ the highest-value food type in their visited patches. The two asocial patches contained two food types, both with nutritional values of +1. Thus, demonstrator choices were arranged in such a way that copying the food patch choice of the best demonstrator would result in the most profitable food patches being found, and copying the exact food type preferences of demonstrators would result in the most nutritious foods being collected and poisonous foods being avoided. Here, we report findings for participant food patch choice only. For further details and analyses on specific food type preferences, please refer to the Supplementary Material (Tables S6-S7; Figure S3).

##### Data collection and preparation

Each time the player or one of the demonstrators entered a food patch, the location and time of the visit were logged. A player was considered to have copied a demonstrator’s food patch choice if they entered a food patch within 20 seconds before or after that demonstrator had entered the same food patch. This time window gave the player enough time to follow a distant demonstrator into a food patch and gave the demonstrator enough time to catch up if the player had overtaken them *en route* to the food patch. If a player entered a food patch that was not visited by any demonstrators within this time window, this was considered an asocial food patch choice. For each participant, the total number of times each demonstrator was copied and the total number of times an asocial decision was made were calculated.

### Post-study questionnaire

After the experiment, participants completed a post-study questionnaire, used to gather information about individual characteristics that may influence social information use, including age and gender. As prior research has indicated that aggression may play a role in social learning (Bandura *et al*., 1961; Easter *et al*., 2022), we also obtained self-assessed aggression scores for each participant, using Bryant and Smith’s (2001) aggression questionnaire, to investigate whether aggression levels influenced social information use in our study. We also considered that a participant’s tendency to play video games may influence their general behaviour within VERSE – e.g. those who play video games often may have a tendency to be more exploratory. Participants were therefore asked how often they played video games in their everyday lives (on a 5-point Likert scale from ‘never’ to ‘daily’) and whether they found the game used in the experiment easy to control (yes/no).

### Statistical analysis

We tested a number of hypotheses concerning participants’ use of social information within our environments, as detailed below. All statistical analyses were conducted in R (v.4.0.4). Due to the number of tests conducted, we report p-values that are corrected for multiple comparisons using the false discovery rate, implemented using the p.adjust() function in R.

### H1: Individuals show a preference towards learning socially

Firstly, we used binomial GLMs (logit link function) to model the tendency of participants to learn socially (copy any demonstrator) or asocially (copy no demonstrator). This was modelled in two ways – first, as the tendency of participants to favour copying a single demonstrator over making an alternative, asocial decision in the *Soc/Asoc* condition and, second, as the tendency of participants to favour copying either demonstrator over making an independent choice across all demonstrator conditions. The latter was modelled for the Route Choice and Foraging tasks only, as the Container task permitted only binary choices that, when two demonstrators were present in the environment, did not allow participants to make decisions completely independently of demonstrator choice.

### H2: Social learning improves performance on tasks

The following analysis was conducted only for the Route Choice and Foraging tasks, because the Container task did not include an asocial option. For each task, we split participants into two groups – those who tended to learn socially (i.e. copied a demonstrator >50% of the time) and those who tended to learn asocially (i.e. copied a demonstrator <50% of the time) across all demonstrator conditions. We then compared the average success (measured as final food scores or final remaining energy values) of largely social versus largely asocial learners across all demonstrator conditions. Since scores could not always be transformed into a Normal distribution, Mann-Whitney U tests were used to determine whether one group was significantly more successful than the other.

### H3: Social learning is biased towards certain types of demonstrators

Binomial GLMs were used to model likelihood that participants in the Same Rewards group copied one demonstrator treatment over the other in the *Dom/Sub, Three/One, Male/Female* and *Large/Small* demonstrator conditions. Since all demonstrators in the Same Rewards group displayed equally profitable behaviours, any biases towards copying particular demonstrators could not be due to differences in demonstrator success.

### H4: Individuals preferentially copy more successful demonstrators

Finally, we investigated whether participants were more likely to copy more successful demonstrators, beyond any innate preferences for those with particular characteristics. To do this, we combined data from the Same Rewards and Different Rewards groups and used binomial GLMs to model the tendency of participants to copy demonstrator A over demonstrator B across the *Dom/Sub, Three/One, Male/Female* and *Large/Small* conditions, with reward group as a predictor. In other words, we analysed whether participants copied demonstrator A significantly more when it displayed more profitable behaviours than demonstrator B, compared to when both demonstrators used equally profitable behaviours. An ANOVA test was then used to establish whether the model that included reward group as a predictor provided a better fit to the data than a corresponding null model.

For hypotheses 1, 3 and 4, we additionally tested whether participants’ individual characteristics (age, gender, aggression, etc) influenced their choices by running a series of binomial GLMs for each hypothesis, as described in the main analysis but with each individual characteristic added as predictors. The results of these additional analyses revealed no consistent or clear pattern. The results and further details on the analysis of individual characteristics are therefore reported in the Supplementary Material (Tables S11-S13).

## Results

### H1: Individuals show a preference towards learning socially

In the *Soc/Asoc* demonstrator condition, where a single AI demonstrated one option and all other options remained undemonstrated, participants in the Same Rewards group were equally likely to learn socially as asocially in the Container (50.0% of all learning events social; z<0.001, p>0.999) and Route Choice (47.1% social, z=-0.840, p=0.446) tasks, but were significantly less likely to learn socially than asocially in the Foraging task (21.4%, z=-9.439, p<0.001) (Figure 2A). This result held both in their initial responses and overall across all choices made (Table 2). In treatment conditions where multiple options were demonstrated by different AIs, participants still had the option to ignore all social information in the Route Choice and Foraging (but not the Container) tasks. Taken across all demonstrator conditions, social learning accounted for significantly fewer learning events than asocial learning in the Route Choice task (41.6%, z=-5.367, p<0.001) and the Foraging task (33.2%, z=-12.900, p<0.001). Individual participants did, however, vary substantially in their tendency to copy and generally used at least some degree of partial copying (see Supplementary Material for discussion, Figure S1).

**Table 2.** Results from binomial GLMs modelling the likelihood of participants copying the choices of demonstrator A over demonstrator B when learning socially, across only the demonstrator conditions where two demonstrators were available, during each of the three tasks. A significant positive intercept term indicates that the participants were more likely to copy demonstrator A, while a significant negative term indicates that participants were more likely copy demonstrator B. Analyses were conducted using data on (upper section) all choices made and (lower) only the initial choices made during each demonstrator condition. Significant p values (< 0.05) are highlighted in bold.

|  |  | Container |  |  |  | Route Choice |  |  |  | Foraging |  |  |  |
| --- | --- | --- | --- | --- | --- | --- | --- | --- | --- | --- | --- | --- | --- |
|  | Condition | Intercept | Std. Error | Z | p | Intercept | Std. Error | z | p | Intercept | Std. Error | z | p |
| All choices | **Dom/Sub** | 0.166 | 0.076 | 2.190 | 0.077 | 0.381 | 0.245 | 1.556 | 0.227 | 0.511 | 0.220 | 2.320 | 0.069 |
|  | **Three/One** | 1.046 | 0.086 | 12.140 | **<0.001** | 1.235 | 0.284 | 4.347 | **<0.001** | 0.490 | 0.181 | 2.702 | **0.034** |
|  | Male/Female | 0.006 | 0.076 | 0.076 | 0.981 | -0.249 | 0.214 | -1.163 | 0.420 | 0.497 | 0.189 | 2.631 | **0.036** |
|  | Large/Small | -0.023 | 0.076 | -0.302 | 0.922 | 0.058 | 0.197 | 0.296 | 0.922 | -0.019 | 0.195 | -0.098 | 0.981 |
| Initial choices | Dom/Sub | 0.465 | 0.246 | 1.895 | 0.139 | 0.647 | 0.372 | 1.737 | 0.178 | 0.916 | 0.418 | 2.190 | 0.077 |
|  | **Three/One** | 0.715 | 0.255 | 2.808 | **0.030** | 2.526 | 0.735 | 3.437 | **0.005** | 0.201 | 0.318 | 0.631 | 0.745 |
|  | Male/Female | -0.114 | 0.239 | -0.478 | 0.844 | -0.636 | 0.412 | -1.543 | 0.227 | 0 | 0.324 | 0 | 1 |
|  | Large/Small | -0.057 | 0.239 | -0.239 | 0.927 | 0.348 | 0.377 | 0.924 | 0.570 | 0.236 | 0.345 | 0.684 | 0.741 |

H3: Social learning is biased towards certain types of demonstrators

**Figure 2.**
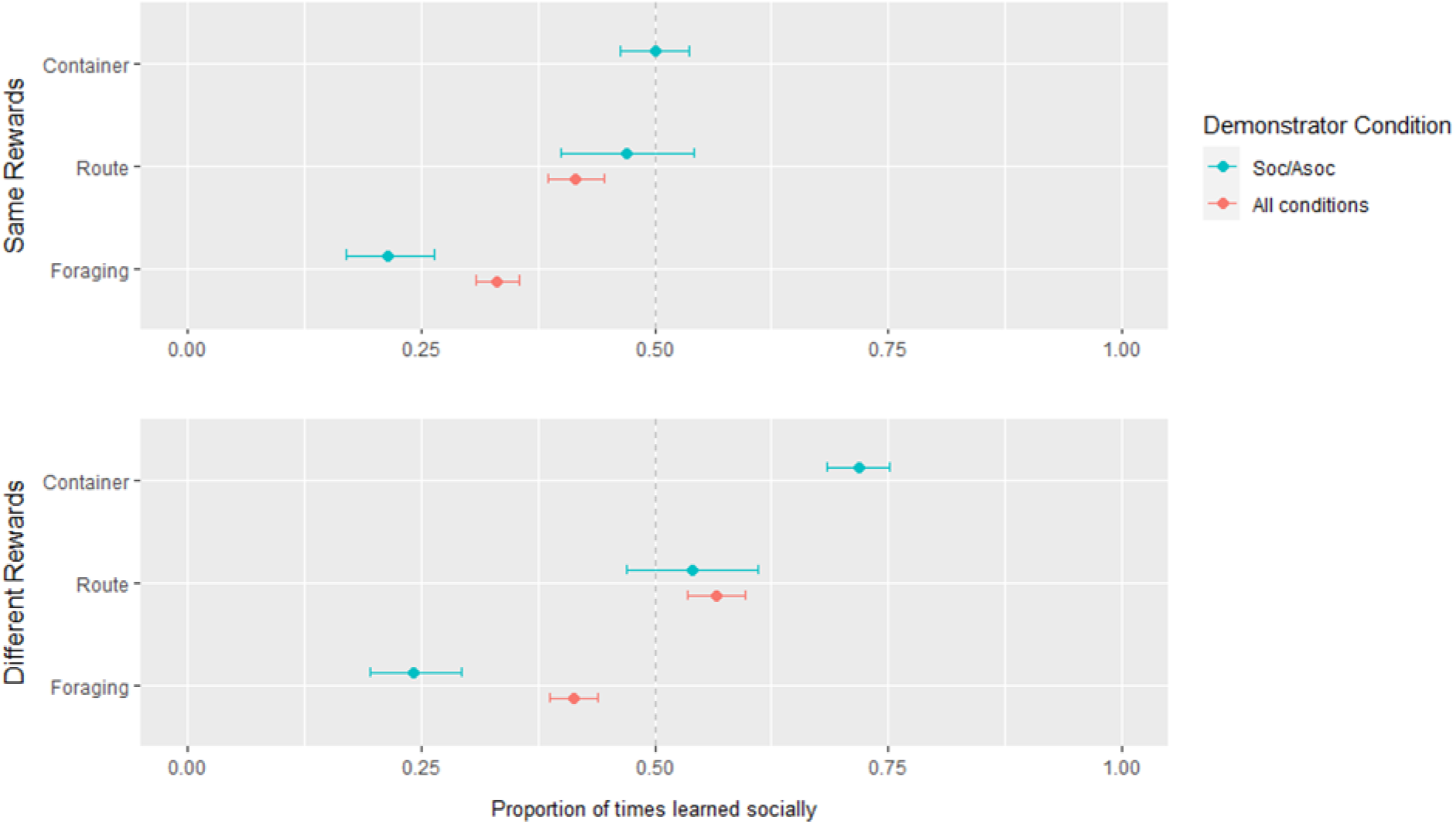
The proportion of times participants learned socially (rather than asocially) during the Container, Route Choice and Foraging tasks, in the *Soc/Asoc* condition (where only one option was demonstrated) and across all demonstrator conditions (where either one or two options were demonstrated). Social learning is defined as copying a demonstrator, while asocial learning is defined as choosing another, undemonstrated option. Results are shown for participants in scenarios where, **(A)** all demonstrated options received equal rewards, and **(B)** demonstrated options received different rewards. Error bars represent Clopper-Pearson 95% confidence intervals. Vertical, dashed reference line is at 0.5 and indicates equal amounts of social and asocial learning occurred.

Compared to scenarios where demonstrated rewards were equal (Same Rewards group), participants were more likely to use social information when rewards varied (Different Rewards group) in the *Soc/Asoc* condition – although this was difference statistically significant in the Container task only (Container: z=8.292, p<0.001; Route Choice: z=1.428, p=0.191; Foraging: z=0.829, p=0.452) – and in general across all demonstrator conditions (Route Choice: z=6.790, p<0.001; Foraging: z=4.550, p<0.001). This resulted in a general preference towards social learning in the Container (71.8% social in the *Soc/Asoc* condition, z=11.15, p<0.001) and Route Choice (54.1% social in the *Soc/Asoc* condition, z=1.18, p=0.238; 56.6% across all demonstrator conditions, z=4.233, p<0.001) tasks, although a preference for asocial learning still remained in the Foraging task (24.1% social in the *Soc/Asoc* condition, z=-8.612, p<0.001; 41.3% across all demonstrator conditions, z=-6.494, p<0.001).

### H2: Social learning improves performance on tasks

When summed across all demonstrator conditions, participants in the Same Rewards group who favoured social learning, on average, completed the Route Choice task with significantly less energy remaining (social score: 73.3%; asocial score: 76.7%; Mann-Whitney U test; *U*(1) = 191, *p* < 0.001) and collected significantly fewer food items in the Foraging tasks (social score: 35.0%; asocial score: 41.2%; Mann-Whitney U test; *U*(1) = 213, *p* = 0.042) than those who favoured asocial learning (Figure 3) (see also Supplementary Material, Figure S2, for results taken from the *Soc/Asoc* condition only). In other words, asocial learning appeared to be more profitable than social learning. However, the ‘adaptive’ value of asocial learning was lower when rewards (and demonstrator success) varied – in the Different Rewards group, participants tended to be equally successful regardless of whether they favoured social or asocial learning in both the Route Choice (social score: 74.5%; asocial score: 74.7%; Mann-Whitney U test; *U*(1) = 498, *p*= 0.402) and Foraging tasks (social: 43.4%; asocial: 40.2%; Mann-Whitney U test; *U*(1) = 652, *p* = 0.221) (Figure 5.5C-D). This result coincided with the general drop in reliance on asocial learning, described above.

**Figure 3.**
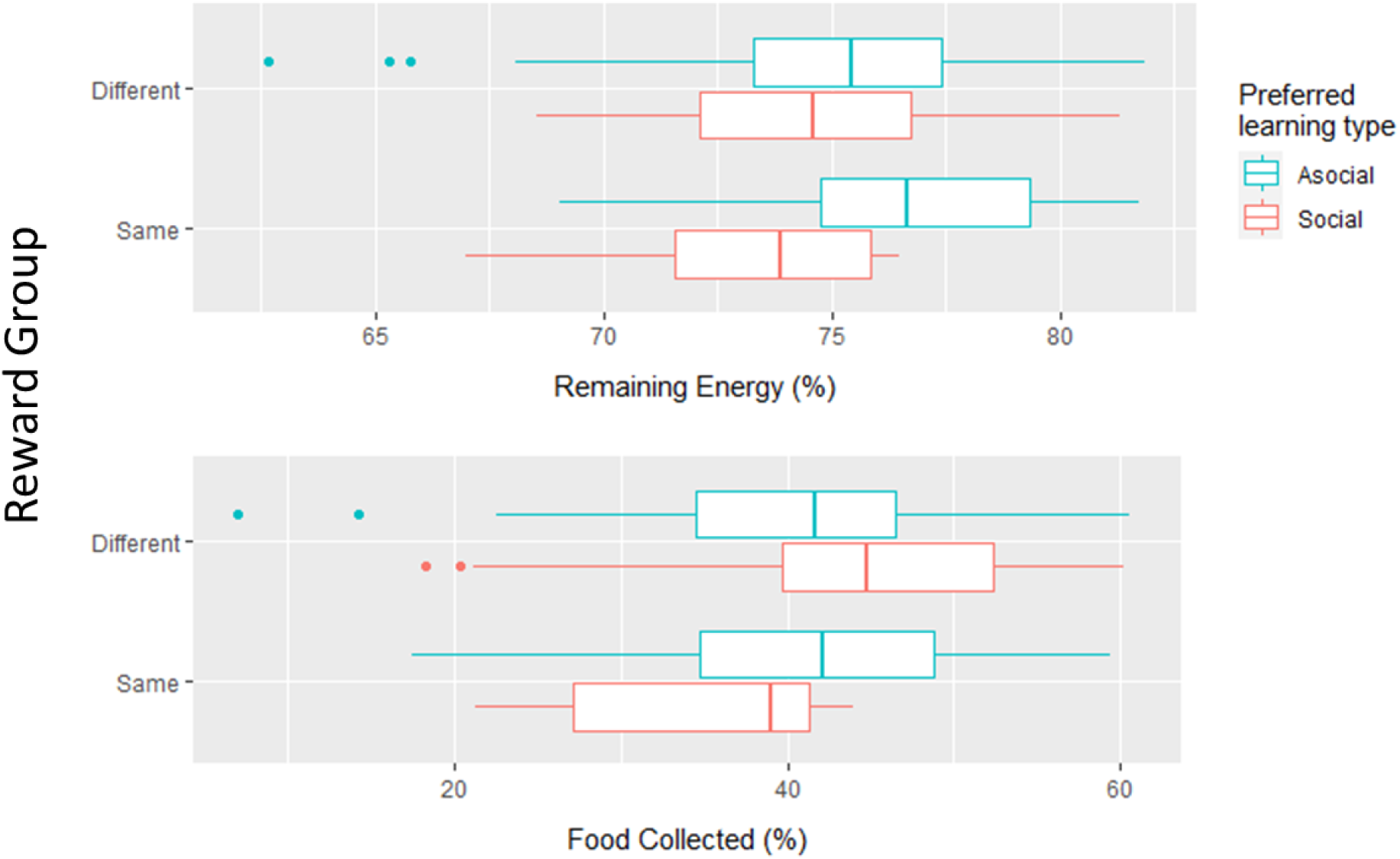
The average success rates of participants who favoured social and asocial learning, on average across all demonstrator conditions, in scenarios where demonstrators received different or the same rewards for their choices, across two different tasks. **(A)** In the Route Choice task, success is measured by the average amount of energy remaining when the end point is reached. **(B)** In the Foraging task, success is measured by the average amount of food collected. Boxplots represent the median and interquartile range. Whiskers extend to 1.5x the interquartile range.

### H3: Social learning is biased towards certain types of demonstrators

#### Dominance bias

When both dominant and subordinate demonstrators were equally successful, participants copied the dominant demonstrator more often than the subordinate in the Container (54.1% of social learning events copied dominant), Route Choice (59.4% dominant) and Foraging (62.5% dominant) tasks (Figure 4A). This bias was statistically significant in the Container and Foraging tasks, but not for the Route Choice task (Table 2). Participants also demonstrated a significant initial bias toward copying the dominant demonstrator in the Foraging task (Table 2).

**Figure 4.**
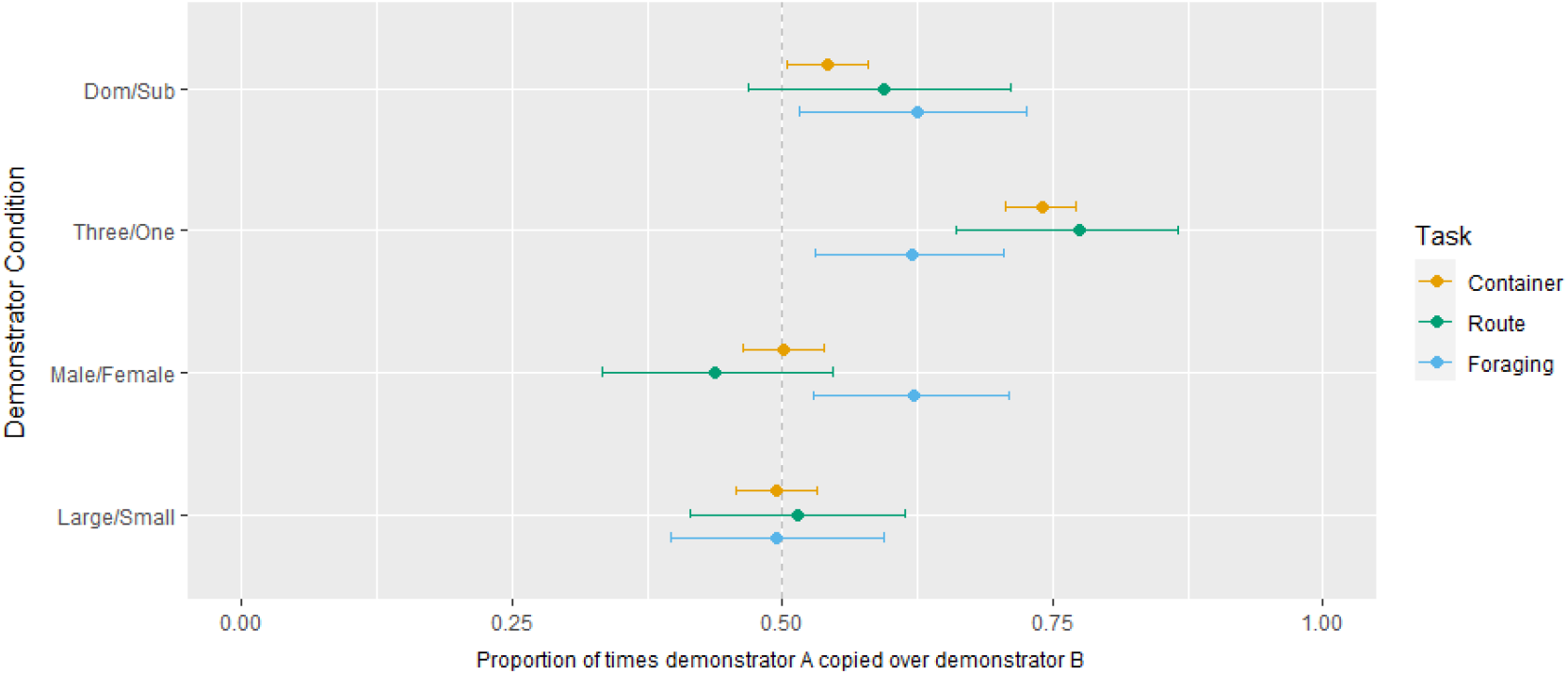
The proportion of times participants copied demonstrator A over demonstrator B in each demonstrator condition – i.e. the dominant over the subordinate demonstrator, the majority (three demonstrators) over the minority (one demonstrator), the male over the female demonstrator and the large over the small demonstrator – when completing each of three tasks. Results are from the Same Rewards group, i.e. in scenarios where each demonstrator displayed equally profitable behaviours. Error bars represent Clopper-Pearson 95% confidence intervals. Vertical, dashed reference line is at 0.5 and indicates no preference for either demonstrator.

#### Frequency bias

Participants copied the majority significantly more often than the minority in all three tasks (Container: 74.0% majority; Route Choice: 77.5% majority; Foraging: 62.0% majority) (Table 2) (Figure 4B). Participants also displayed a significant initial bias towards copying the majority in the Container and Route Choice tasks (Table 2).

#### Sex bias

Participants did not display any sex-based bias in the Container task (50.1% male). The female demonstrator was copied more often in the Route Choice task (56.2% female) but this was not statistically significant (Table 2). The male demonstrator, however, was copied significantly more often in the Foraging task (62.2% male) (Table 2) (Figure 4C).

#### Size bias

Participants did not display any significant bias towards demonstrators of different sizes in any of the tasks (Container: 49.4% large; Route Choice: 51.5% large; Foraging: 49.5% large) (Table 2) (Figure 4D).

### H4: Individuals preferentially copy more successful demonstrators

GLMs predicting participants’ likelihood of copying demonstrator A over B performed significantly better when reward group was included as a factor (Supplementary Material, Table S8), suggesting differences in social information use when there was variable success between demonstrators. Specifically, when learning socially, participants in the Different Rewards group were significantly more likely than participants in the Same Rewards group to copy demonstrator A (the more successful demonstrator) than demonstrator B (the less successful) in the Container (Different Rewards: 75.7% of social learning events copied demonstrator A, Same Rewards: 56.9%, z=11.427, p<0.001), Route Choice (Different Rewards: 74.2% copied A, Same Rewards: 56.6%, z=5.163, p<0.001) and Foraging (Different Rewards: 72.8% copied A, Same Rewards 59.2%, z=4.400, p<0.001) tasks (Figure 5). In other words, participants were significantly more likely to copy demonstrator A over B when demonstrator A displayed more successful behaviours. This pattern was consistent across almost all demonstrator condition/task combinations (Supplementary Material, Table S9), suggesting that a preference for copying the most successful demonstrators was not influenced by context or the specific demonstrators involved.

**Figure 5.**
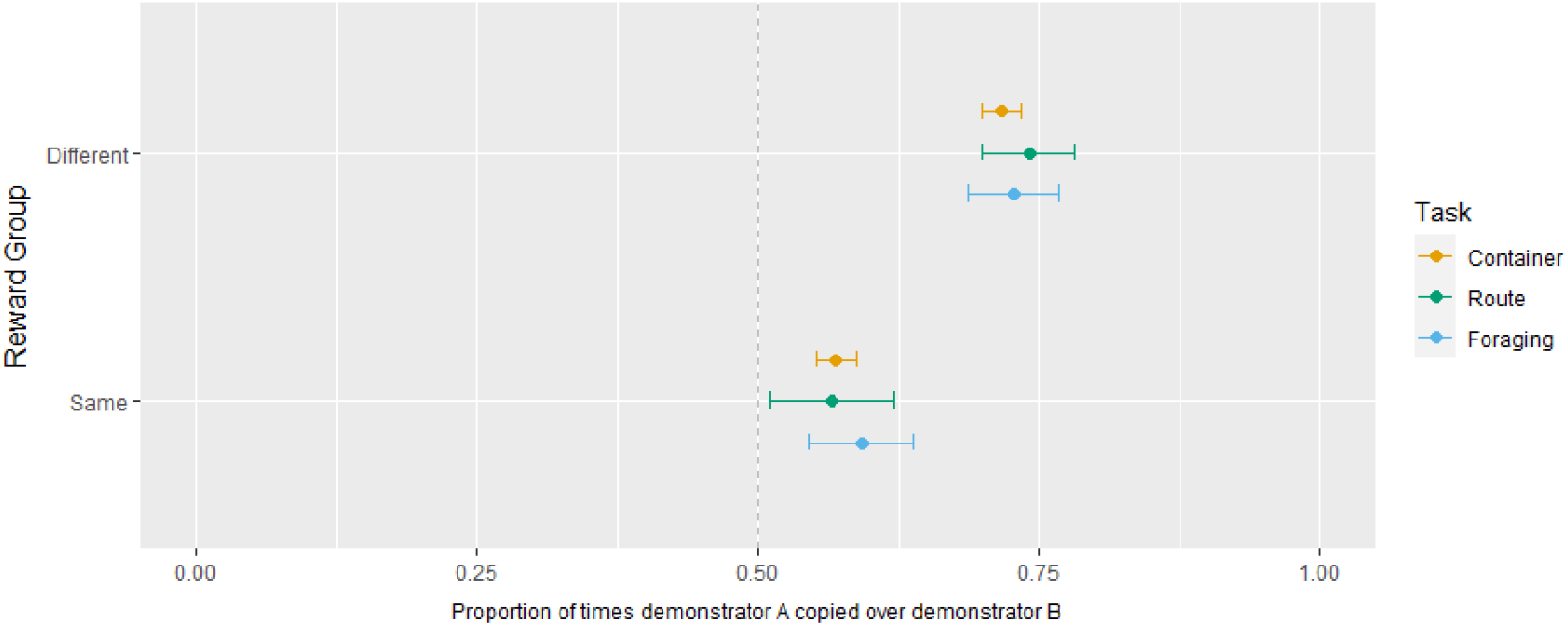
The proportion of times participants copied the most successful demonstrator (demonstrator A) over the least successful (demonstrator B) in the Different and Same Rewards group (i.e. in scenarios where demonstrated rewards either varied or were equal or varied), across the three tasks. A higher proportion copying demonstrator A when demonstrator rewards varied compared to when they were equal indicates a bias for copying successful demonstrators, beyond any innate preferences for demonstrators with particular characteristics. Data is collated across the *Dom/Sub, Three/One, Male/Female* and *Large/Small* demonstrator conditions, i.e. conditions in which two demonstrators were present. Error bars represent Clopper-Pearson 95% confidence intervals. Vertical, dashed reference line is at 0.5 and indicates no preference for either demonstrator.

## Discussion

We have demonstrated a novel approach to studying human social learning behaviour, through the use of specially designed, three-dimensional virtual environments that allow participants to express more natural behaviours, across more realistic scenarios and spatial scales, than is often possible in laboratory conditions. Using this technology, we found that participants generally preferred to learn asocially (although this varied across contexts) and that, in certain environments, asocial learning allowed participants to exploit more profitable, undemonstrated solutions. Participants were strongly biased towards copying the majority over the minority and towards copying more successful demonstrators. Below, we discuss our findings in detail, including how they compare to previous research from more restrictive laboratory experiments and evidence for similarities in social information use across other, unrelated taxa.

Perhaps our most striking finding concerned the way in which participants used social versus asocial information. Namely, participants displayed a general preference for learning a task independently rather than learning socially. The fact that humans are highly cultural creatures – and culture is, by definition, the product of social transmission of behaviours through a population (Laland and Hoppitt, 2003) – suggests that our decisions are highly influenced by the decisions of those around us. Theoretical analyses also agree that social learning, provided it is strategic, tends to outcompete asocial learning (Rendell *et al*., 2010; Boyd and Richerson, 1985). Despite this, several empirical studies have demonstrated, in agreement with our own findings, that adult humans are often biased towards using information they have gained independently (Yaniv and Kleinberger, 2000; Eriksson and Strimling, 2009; Weizsäcker, 2010). This has previously been attributed to the relatively simple, two-option tasks used in such studies, which allow participants to be confident enough in their own decisions to avoid the use of social information (Morgan *et al*., 2012; Muthukrishna *et al*., 2016). On the contrary, our results suggest that a bias for learning independently also exists when completing complex, multi-option tasks within immersive, spatially explicit environments.

We found little evidence to suggest that the individual characteristics of participants (namely age, gender and aggression levels) influenced social learning decisions – although it is worth emphasising the narrow age range of our study group, which may not be representative of all communities. It is, however, possible that some other, untested traits (e.g. ethnicity, education, motivation, prior knowledge) correlated with social information use in our study group. Instead, participants’ tendency to learn socially varied in several ways depending on the scenario. Participants displayed an increased reliance on asocial learning as the tasks became more complex. This is somewhat surprising – one might expect that, when faced with more complex tasks, it would become more difficult to find solutions independently (e.g. require more random searches of the environment) and so informed demonstrators would become more valuable sources of information. Previous work tends to agree that both human and non-human animals are more prone to copying others when tasks are more difficult (Baron *et al*., 1996; Laland, 2004; Kendal, Kendal, *et al*., 2009; Morgan *et al*., 2012) – although this is not the case for all species (e.g. task difficulty does not appear to influence social information use in otters; Ladds *et al*., 2017). Nevertheless, in scenarios where demonstrators displayed equally profitable behaviours, we found that participants who were prone to learning asocially tended to be more successful, suggesting that a reliance on asocial learning was beneficial even when tasks were complex. We therefore suggest that this pattern of behaviour may not be due to task complexity *per se*, but due to a combination of greater opportunities for asocial learning and increased difficulty in acquiring social information due to spatial factors as task complexity increased.

The tasks used in our study essentially formed a continuum in both spatial scale and the variety of alternative, undemonstrated solutions that could be explored from Container task (binary choice, small scale), to Route Choice task (defined paths, option for independent decisions, large scale), to the Foraging task (complex search patterns across larger landscapes, demonstrators are scattered, high level of flexibility in solutions). Muthukrishna and colleagues (2016) have previously demonstrated that people are more likely to change their decisions in favour of demonstrated solutions as the number of possible options increases, which at first glance appears to contradict our results. However, Muthukrishna and colleagues presented social information for all options, thus there were no options that could be discovered via purely independent exploration. We have, in contrast, demonstrated that participants will independently explore complex environments in search of more profitable alternative solutions (e.g. shorter routes or undiscovered food patches) than are currently available in the social information pool – which has clear adaptive value if it prevents individuals from becoming fixed on less profitable behaviours by constant copying. In addition, from the Container to the Foraging task, demonstrators became more scattered across larger spatial scales, which may have effectively increased the cost of acquiring social information by making it more difficult to determine what everyone is doing at a particular time, thus increasing participants’ reliance on asocial learning within larger environments. Social learning is often assumed to be less costly than asocial learning, however an increase in the cost of acquiring social information reduces its adaptive value (Mesoudi, 2008). Taken together, this would explain the increased reliance on asocial learning as task and environmental complexity increased. Due to the restrictions of laboratory conditions that more closely resemble our Container task, previous research has not fully considered the influence of spatial asynchrony of demonstrators on a participant’s ability to use social information and we suggest that this may lead to overestimations of human reliance on social information in more realistic scenarios.

We also reported an increase in social learning when rewards and demonstrator success varied compared to scenarios where demonstrated behaviours received equal payoffs. Participants who relied more heavily on social learning were no more successful than largely asocial learners in these scenarios and so there appeared to be no direct benefit to this increase in social learning. Therefore, it may be that participants were instead responding to greater uncertainty about their own independent choices caused by variability in potential rewards. This result is indicative of a number of potential social learning strategies, including *copy-when-uncertain, copy-if-dissatisfied* and – since asocial learning was less profitable and therefore more costly to acquire in terms of time and energy when the environment contained varying payoffs – *copy-when-asocial-learning-is-costly* strategies (Laland, 2004; Boyd and Richerson, 1985). Theoretical analyses predict that individuals will increase their reliance on social information, which is cheaper but less reliable than sampling directly from the environment, when asocial learning is too costly to acquire or when it is difficult to determine the best behaviour to use by oneself – e.g. due to variation in yield (Boyd and Richerson, 1985) or delayed rewards (Caldwell and Eve, 2014) – which appears consistent with the results reported here. This pattern of social information use is similar to that of starlings and sticklebacks, both of which prefer to tackle foraging problems independently, but will weight their use of social and asocial information depending on their reliability and how difficult they are to acquire (Templeton and Giraldeau, 1996; van Bergen *et al*., 2004).

We found little evidence to suggest that the demonstrators’ physical and/or behavioural characteristics influenced who participants learned from – apart from a weak, nonsignificant bias towards copying dominant over subordinate demonstrators. A tendency to copy dominant individuals has been demonstrated in some species, most notably domestic hens (*Gallus gallus domesticus*) (Nicol and Pope, 1994; Nicol and Pope, 1999) and chimpanzees (*Pan troglodytes*) (Kendal *et al*., 2015); however, in some cases it is unclear whether this is due to a genuine observer bias or whether dominant individuals simply restrict subordinates’ access to novel task, thus making them less effective as demonstrators (Watson *et al*., 2017). A study by Wood *et al*. (2013) demonstrated that human children selectively watch more dominant demonstrators in an open diffusion experiment, even when dominant individuals do not monopolise the novel task. The influence of dominance over human decisions may therefore warrant some further study, perhaps using a more complex dominance hierarchy than employed in our study.

Compared to individual demonstrator characteristics, the frequency of demonstrators using particular options appeared to have a much stronger influence on participants’ social information use. When participants chose to use social information rather than attempt a task independently, there was strong evidence of an innate bias towards copying the majority, even when it was no more profitable to do so. This pattern occurred across all tasks and is in agreement with previous, lab-based human experiments (Morgan *et al*., 2012; Haun *et al*., 2012; Muthukrishna *et al*., 2016; Deffner *et al*., 2020), suggesting that humans possess a majority bias in a wide range of contexts and task complexities. An innate tendency to copy the majority (also known as a ‘positive frequency-dependent’ bias) is likely to be an adaptive strategy when learning to survive in a novel environment because, if many individuals already use a specific behaviour within that environment, it is likely to be an adaptive behaviour that receives adequate payoff to sustain those individuals. In addition, such a strategy is fairly easy to implement as it does not require different demonstrators to be assessed based on their individual characteristics or success rates. It is therefore unsurprising that a bias toward copying the majority strategy has been identified across a number of taxa, from humans and other primates to birds to fish (Morgan *et al*., 2012; Van de Waal *et al*., 2013; Aplin *et al*, 2015; Day *et al*., 2001) and that theoretical analyses agree on its adaptive value (Boyd and Richerson, 1985).

When we introduced uneven payoffs to different demonstrated strategies, participants imitated the successful demonstrators and showed an increase in reliance on social over asocial information, as discussed above. Payoff-based social learning has been reported in human lab studies (Mesoudi and O’Brien, 2008; Mesoudi, 2011; Atkisson *et al*., 2012; Molleman *et al*., 2014). However, those studies tended to use comparatively simple sources of social information that do not incorporate the potential difficulties in assessing multiple behaviours being performed by multiple demonstrators asynchronously in both space and time. Copying the most successful individuals has obvious adaptive benefits, as confirmed by theoretical analyses (Schlag, 1998; Grove, 2018); however, gaining this information in real-world situations comes with the time and energy costs of tracking and comparing the success rates of multiple individuals for a given task, which is likely to become more difficult across larger spatial scales. Nine-spined sticklebacks use public information to assess the profitability of food patches and appear to deploy a ‘hill-climbing’ strategy when deciding where to feed, favouring the choices of individuals more successful than themselves until an optimal behaviour is reached (Coolen *et al*., 2003; Kendal, Rendell, *et al*., 2009). Similarly, field experiments on wild white-faced capuchins (*Cebus capucinus*) and vervet monkeys show that both species preferentially learn the food extraction techniques of conspecifics who receive the highest yield (Barrett *et al*., 2017; Canteloup *et al*., 2021). This strategic use of social information essentially allows individuals to reap the benefits associated with both social and asocial learning – reducing the time and risk associated with independent learning while ensuring that social information is accurate and profitable.

## Summary and conclusions

We have demonstrated a novel approach to studying human social learning behaviour, through the use of specially designed, three-dimensional virtual environments, developed using gaming technology, that allows participants to express more natural behaviours, across more realistic spatial scales, than is often possible in laboratory conditions. Using this technology, we explored human social learning strategies in a series of experiments inspired by the animal literature – thus allowing direct generalisations to be made across the human/non-human divide. In general, participants displayed an innate bias towards learning independently rather than socially in these complex environments. While we found little evidence to suggest that participants were biased toward copying demonstrators with particular individual characteristics, we did find clear evidence for two social learning biases previously highlighted in both the human and animal literature – namely a tendency to copy the majority and a tendency to copy successful individuals. These happen to be the two of the most extensively studied social learning strategies, demonstrated before both in humans and across other taxa, and there is evidence to suggest that both strategies are adaptive ways of gaining the most profitable social information. Overall, our novel approach to studying human social learning allowed us to study how people learn in novel, realistic environments, taking into account the real-life complexities of gathering information both socially and asocially. We encourage future researchers to focus on testing these foundations against more realistic scenarios in order to better understand the evolutionary importance of the way in which we learn from one another.

## Supporting information

Supplementary Material, containing additional methodological details and additional analyses

## Acknowledgements

A big thanks to Will Hoppitt for your help conceptualising the experiment in the early stages and also for your statistical advice.

## Data accessibility

All raw data, organised datasets and R code required to reproduce the results of this study are available in the following FigShare repository: https://doi.org/10.6084/m9.figshare.19196600. Additional supplementary analyses and results are provided in the Supplementary Material.

## Ethical approval

Ethical approval for this study was obtained by the Faculty of Biological Sciences Research Ethics Committee, University of Leeds (LTSBIO-029).

