## Supplementary Material, containing additional methodological details and additional analyses for "Human social learning biases in immersive virtual environments"

### 1.1. Supplementary Material

#### 1.1.1. Presentation order of tasks and demonstrator conditions across participants

Both the order of task presentation and the order of demonstrator condition presentation were randomised across participants to reduce the impact of presentation order on social information use (Tables S1, S2). The order of presentation for both the tasks and the demonstrator conditions within each task followed a balanced Latin square design where possible. For the Route Choice and Foraging tasks, where several replicates were produced for each demonstrator condition, each with a distinctly different environment, the replicates used were also randomised across participants (Table S3).

**Table S1.** The randomised order of tasks and demonstrator conditions for each participant. Demonstrator orders are represented by letters, which correspond to the appropriate rows in Table S2.

| | Task 1 | | Task 2 | | Task 3 | |
|---|---|---|---|---|---|---|
| Participant | Task | Demonstrator order | Task | Demonstrator order | Task | Demonstrator order |
| 1 | Route finding | A | Tokens | B | Foraging | F |
| 2 | Route finding | B | Tokens | C | Foraging | A |
| 3 | Route finding | C | Tokens | D | Foraging | B |
| 4 | Route finding | D | Tokens | E | Foraging | C |
| 5 | Route finding | E | Tokens | F | Foraging | D |
| 6 | Route finding | F | Tokens | A | Foraging | E |
| 7 | Route finding | A | Foraging | B | Tokens | F |
| 8 | Route finding | B | Foraging | C | Tokens | A |
| 9 | Route finding | C | Foraging | D | Tokens | B |
| 10 | Route finding | D | Foraging | E | Tokens | C |
| 11 | Route finding | E | Foraging | F | Tokens | D |
| 12 | Route finding | F | Foraging | A | Tokens | E |
| 13 | Tokens | A | Foraging | B | Route finding | F |
| 14 | Tokens | B | Foraging | C | Route finding | A |
| 15 | Tokens | C | Foraging | D | Route finding | B |
| 16 | Tokens | D | Foraging | E | Route finding | C |
| 17 | Tokens | E | Foraging | F | Route finding | D |
| 18 | Tokens | F | Foraging | A | Route finding | E |

| | | | | | | |
|---|---|---|---|---|---|---|
| 19 | Tokens | A | Route finding | B | Foraging | F |
| 20 | Tokens | B | Route finding | C | Foraging | A |
| 21 | Tokens | C | Route finding | D | Foraging | B |
| 22 | Tokens | D | Route finding | E | Foraging | C |
| 23 | Tokens | E | Route finding | F | Foraging | D |
| 24 | Tokens | F | Route finding | A | Foraging | E |
| 25 | Foraging | A | Route finding | B | Tokens | F |
| 26 | Foraging | B | Route finding | C | Tokens | A |
| 27 | Foraging | C | Route finding | D | Tokens | B |
| 28 | Foraging | D | Route finding | E | Tokens | C |
| 29 | Foraging | E | Route finding | F | Tokens | D |
| 30 | Foraging | F | Route finding | A | Tokens | E |
| 31 | Foraging | A | Tokens | B | Route finding | F |
| 32 | Foraging | B | Tokens | C | Route finding | A |
| 33 | Foraging | C | Tokens | D | Route finding | B |
| 34 | Foraging | D | Tokens | E | Route finding | C |
| 35 | Foraging | E | Tokens | F | Route finding | D |
| 36 | Foraging | F | Tokens | A | Route finding | E |

**Table S2.** The six demonstrator condition orders, randomised across participants.

| | Order | | | | | |
|---|---|---|---|---|---|---|
| | **1st** | **2nd** | **3rd** | **4th** | **5th** | **6th** |
| **A** | *Asocial* | *Soc/Asoc* | *Dom/Sub* | *Three/One* | *Male/Female* | *Large/Small* |
| **B** | *Soc/Asoc* | *Three/One* | *Asocial* | *Large/Small* | *Dom/Sub* | *Male/Female* |
| **C** | *Three/One* | *Large/Small* | *Soc/Asoc* | *Male/Female* | *Asocial* | *Dom/Sub* |
| **D** | *Large/Small* | *Male/Female* | *Three/One* | *Dom/Sub* | *Soc/Asoc* | *Asocial* |
| **E** | *Male/Female* | *Dom/Sub* | *Large/Small* | *Asocial* | *Three/One* | *Soc/Asoc* |
| **F** | *Dom/Sub* | *Asocial* | *Male/Female* | *Soc/Asoc* | *Large/Small* | *Three/One* |

**Table S3.** The replicates used and their order of presentation for the Route Choice and Foraging tasks, for each of the demonstrator orders shown in Table S2.

| Demonstrator condition order | Route Choice (three of a possible five) | Foraging (one of a possible three) |
|---|---|---|
| A | 1, 2, 3 | 1 |
| B | 4, 5, 1 | 2 |
| C | 3, 4, 2 | 3 |
| D | 2, 1, 5 | 1 |
| E | 5, 3, 4 | 2 |
| F | 1, 4, 3 | 3 |

#### 1.1.2. Perceptions of the AI models used.

Prior to commencing the experiment, it was important to understand whether people, in general, perceived the AI models in the way intended by the researchers – e.g. viewed the 'female' AI as female, the 'dominant' AI as dominant to the 'subordinate' AI, etc. A group of 41 University of Leeds postgraduate students and academic staff volunteered, in response to an email request, to complete an online questionnaire using Microsoft Forms, in which they gave their perceptions of the AI models to be used in the main experiment. In this questionnaire, participants were presented with four sets of images, each displaying a pair of AI models (Table S4). For each set, the following questions were asked: (i) In your opinion, what gender(s) are the two figures? (ii) In your opinion, does one figure appear to be more dominant than the other? (iii) In your opinion, which figure appears friendlier or more approachable? (iv) In your opinion, which figure appears more aggressive?

**Table S4.** Images presented to a group of participants prior to the main experiment, showing four sets of two AI models. These AI models were to be used in the main experiment to detect biases towards learning from individuals with certain characteristics. This questionnaire was therefore used to establish whether people generally perceived the AIs in the intended way.

| **Set** | **Description** | **Image displayed** |
|---|---|---|

| | | |
|---|---|---|
| 1 | Figure 1 is a small AI, Figure 2 is a large AI. Both figures are the ‘default’ body shape used in the experiment. Other than overall size, all proportions are the same. | 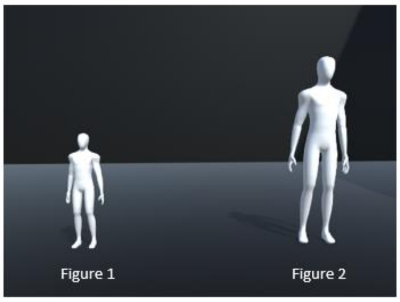<br><br>*Front view*<br>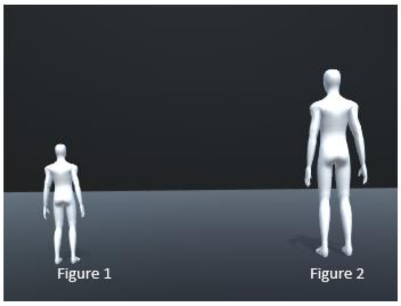<br><br>*Back view* |
| 2 | Figure 1 is a ‘dominant’ AI, which is more muscular and stands upright. Figure 2 is a ‘subordinate’ AI, which is emaciated in appearance and stands hunched. An additional side view was given to make this difference in posture clear. | 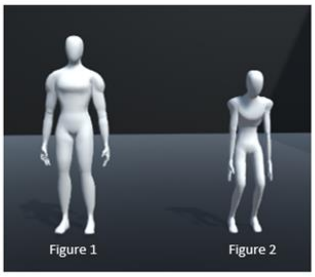<br><br>*Front view*<br>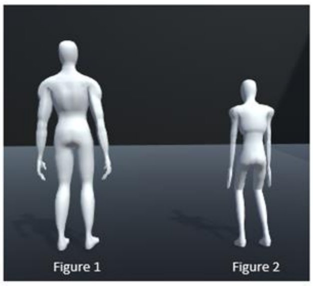<br><br>*Back view*<br>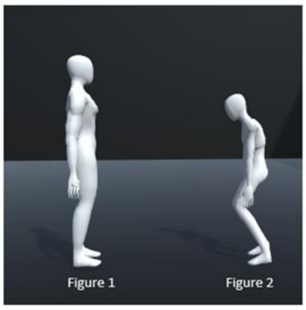<br><br>*Side view* |

| | | |
|---|---|---|
| 3 | Figure 1 is a ‘male’ AI (which was also used as the default AI in the *Three/One* and *Large/Small* demonstrator conditions). Figure 2 is a ‘female’ AI. | 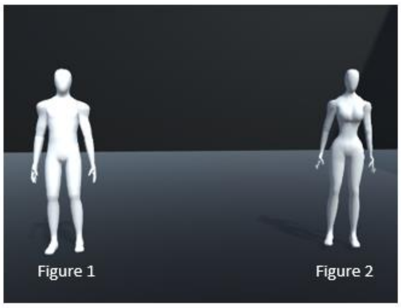 *Front view*<br>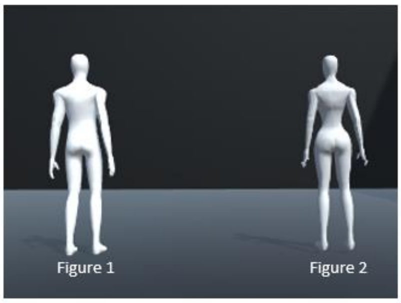 *Back view* |
| 4 | ‘Dominant’ and ‘subordinate’ AIs with interaction. Figures are the same as in set 2, but now display the character interactions that will be displayed at the beginning of each *Dom/Sub* demonstrator condition in the main experiment. | 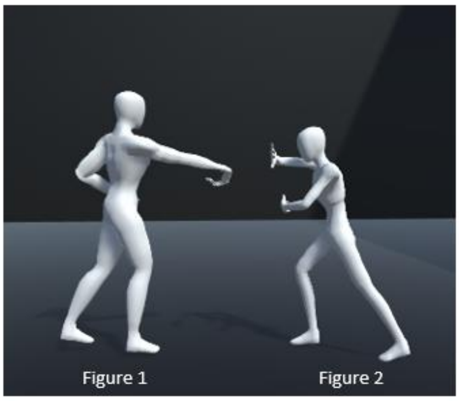 |

The results of the questionnaire are displayed in Table S5. In response to AIs of different sizes, 92% of participants agreed that both AIs were male. Both AIs were viewed as equally dominant by 48% of participants, and the larger AI was viewed as dominant to the smaller AI by 43% of participants. This suggests that any bias towards learning from larger AIs in the main experiment could be, in part, due to a dominance bias. Aggressiveness and approachability of AIs based on their size varied between participants.

In response to a ‘male’ and a ‘female’ AI, 100% of participants agreed that the ‘male’ AI was male and the ‘female’ AI was female. Gender was therefore an easily distinguishable trait and participants perceived AI genders in the intended way. Interestingly, this consensus suggests that participants’ perception of ‘gender’ aligned with stereotypical physical sex differences. In general, both male and female AIs were viewed as equally dominant, approachable and aggressive. Thus, we can be confident that that any learning biases detected in the *Male/Female* demonstrator condition will be due to their perceived gender differences and not due to any of the alternate characteristics considered here. It is also worth noting that the 'male' AI displayed here was the default AI used across demonstrator conditions that did not require physical characteristics to be altered (namely the *Soc/Asoc* and *Three/One* conditions), and so participants will likely view all AIs as male in these conditions as well.

In response to the 'dominant' and 'subordinate' AIs, 95% of participants agreed that the 'dominant' AI was dominant to the 'subordinate' (which dropped slightly to 87% when actions were added), and the 'subordinate' AI was never viewed as dominant to the 'dominant' AI. Thus, participants perceived the dominance status of these AIs in the intended way. When the two AIs were displayed with no actions, there was variation in the genders they were both perceived as, but the majority of participants agreed that the 'subordinate' was female. When actions were included in the image, the 'dominant' AI was viewed as male by 80% of participants and the 'subordinate' AI was viewed as female by 72% of participants. Any dominance-based learning biases detected in the main experiment could therefore be due, at least in part, to perceived differences in their gender. Interestingly, when the AIs were displayed without associated actions, the majority of participants found the 'dominant' AI to be more approachable, despite a general consensus that the 'dominant' AI was either equally or more aggressive-looking than the 'subordinate' AI. When the AIs were shown again with their associated actions, however, perceptions on approachability and aggression were altered, with the 'dominant' AI generally viewed as more aggressive-looking than the 'subordinate' AI, and the majority of participants now perceiving the 'subordinate' AI as more or equally approachable compared to the 'dominant' AI. Perceptions of aggressiveness and, in particular, approachability therefore appear independent of perceptions of dominance – with participants generally finding the AI dominant in appearance alone more approachable. This may be because the hunched, emaciated appearance of the 'subordinate' AI was viewed as intimidating. However, when the AIs displayed dominance-related actions, the relatively aggressive action of the 'dominant' AI made them appear less approachable. As dominance is often directly linked to aggression, including the dominance-related interactions between these two AIs thus seems an important factor to reinforce their perceived relationship.

**Table S5.** Results of the pre-study questionnaire on the perceptions of different AI models, showing participants' perceptions of the gender, dominance, approachability and aggressiveness of the two AIs in each pair. Values represent the proportion of participants expressing a particular opinion about the characteristics of each AI pair.

| | | AI pair | | | |
|---|---|---|---|---|---|
| **Characteristic** | **Opinion** | **Small/Large** | **Male/Female** | **Dom/Sub** | **Dom/Sub (with actions)** |
| Gender | Fig 1 male, fig 2 female. | 0.04 | 1.00 | 0.38 | 0.54 |
| | Fig 1 female, fig 2 male. | 0.04 | 0 | 0.05 | 0.03 |
| | Both male. | 0.91 | 0 | 0.23 | 0.26 |
| | Both female. | 0 | 0 | 0.35 | 0.18 |
| Dominance | Fig 1 is dominant. | 0.09 | 0.03 | 0.95 | 0.87 |
| | Fig 2 is dominant. | 0.43 | 0.08 | 0 | 0 |
| | Both equally dominant. | 0.48 | 0.90 | 0.05 | 0.13 |
| Approachability | Fig 1 is more approachable. | 0.30 | 0.08 | 0.68 | 0.08 |
| | Fig 2 is more approachable. | 0.22 | 0.15 | 0.13 | 0.51 |
| | Both equally approachable. | 0.48 | 0.78 | 0.20 | 0.41 |
| Aggressiveness | Fig 1 is more aggressive. | 0.13 | 0.03 | 0.40 | 0.84 |
| | Fig 2 is more aggressive. | 0.35 | 0.13 | 0.08 | 0.03 |
| | Both equally aggressive. | 0.52 | 0.85 | 0.52 | 0.13 |

#### 1.1.1. Additional analysis: Variation in participants' use of social information

Previous research suggests that some individuals are mainly asocial learners while others are mainly social learners (Efferson *et al.,* 2008; Toelch *et al.,* 2014; Miu *et al.,* 2020). Here, we investigate whether participants in the Same Rewards group varied in their reliance on social learning. Using a series of Spearman's rank-order correlation tests, we also determine whether individual participants were consistent in their use of social information across tasks – i.e. whether certain individuals could be categorised based on their overall reliance on social information across different contexts. Specifically, we obtained pairwise correlation measures for the proportion of times each participant learned socially within the *Soc/Asoc* demonstrator condition for all three tasks, and across all demonstrator conditions for the Route Choice and Foraging tasks.

For each task, participants in the Same Rewards group varied substantially in the proportion of times they chose to learn socially. This ranged from a complete ignorance of to a complete reliance on social information, depending on the task and participant in question. In the

*Soc/Asoc* condition, the proportion of times a participant copied the single AI ranged from 0 to 1 in all three tasks (Container average: 0.5 ± 0.21 SD; Route Choice average: 0.47 ± 0.38 SD; Foraging task average: 0.22 ± 0.29 SD). There was no significant correlation between a participant's tendency to copy versus ignore the single AI across the three tasks in the *Soc/Asoc* condition (Spearman's rank tests: Container—Route, $r_s(65) = -0.02$; $p = 0.87$; Container—Foraging, $r_s(67) = -0.03$; $p = 0.81$; Route—Foraging, $r_s(66) = 0.17$; $p = 0.18$) (Figure S1A-C). When taken across all demonstrator conditions, the proportion of times a participant copied either demonstrator rather than opting for an alternative, asocial option ranged from 0.07 to 1 for the Route Choice task (average: 0.41 ± 0.27 SD) and from 0 to 0.83 for the Foraging task (average: 0.34 ± 0.20 SD). There was a moderate, statistically significant correlation in each participant's tendency to copy rather than find an alternative solution across the Route Choice and Foraging tasks when data was combined across demonstrator conditions (Spearman's rank test: $r_s(67) = 0.30$; $p = 0.01$) (Figure S1D). Note that combining participant choices across all demonstrator conditions may give a more precise measure of social learning tendency due to the larger number of replicates the data is taken over (5 replicates for each participant when taken across all demonstrator conditions, compared to 1 replicate for the *Soc/Asoc* condition only).

Overall, this suggests that participants displayed some consistency in their inclination towards using social information across different tasks and demonstrator conditions (a correlation highly comparable to Toelch *et al.*, 2014), but not to the extent that they could be categorised as solely social or asocial learners (e.g. the 'conformists' and 'mavericks' identified in Efferson *et al.*'s 2008 study).

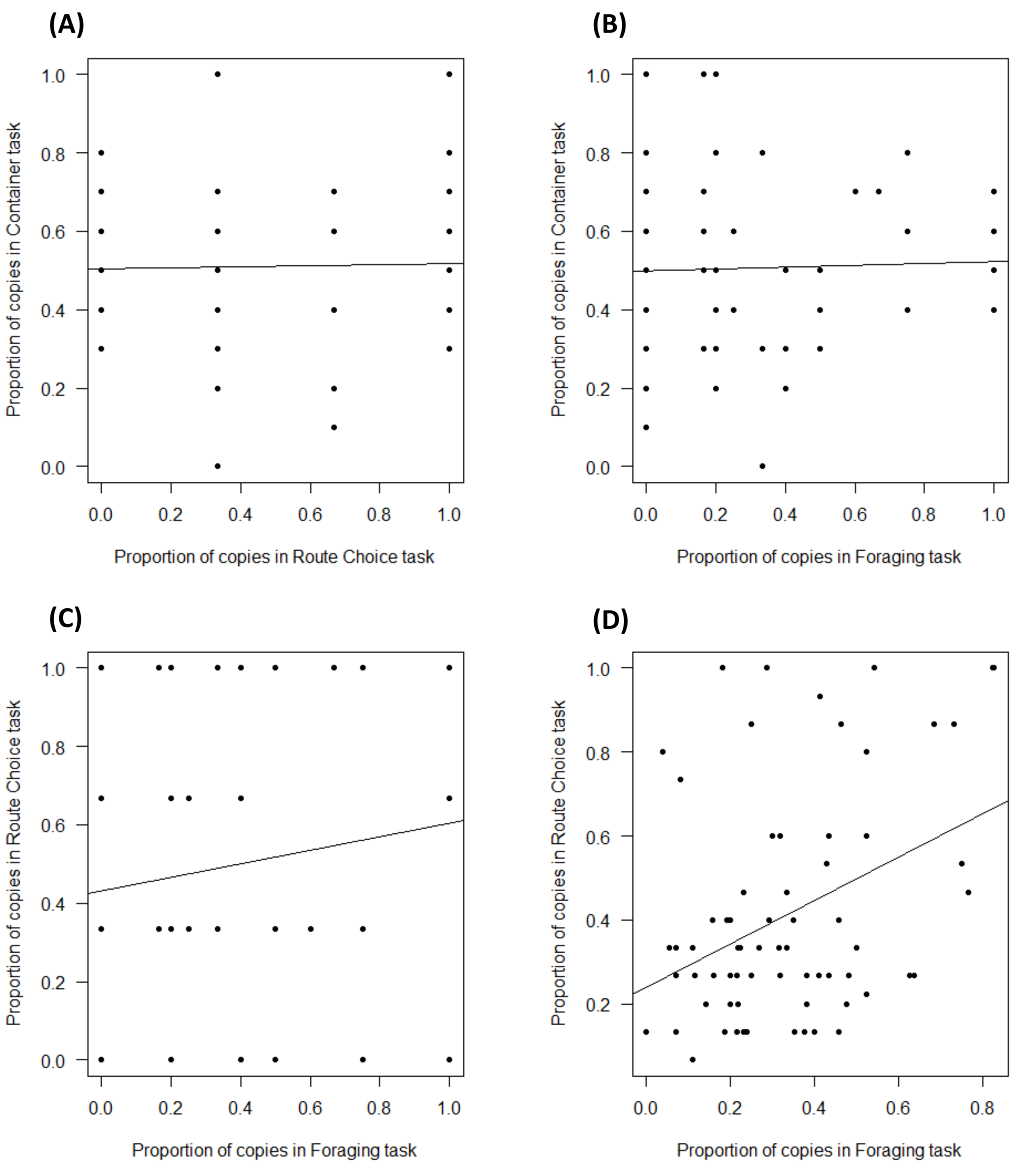

**Figure S1.** Correlations between the proportion of times each participant used social information in each pair of tasks. **(A-C)** For the *Soc/Asoc* condition only (i.e. when only one demonstrator was present and all other choices were considered asocial learning): **(A)** Container ~ Route Choice. **(B)** Container ~ Foraging. **(C)** Route Choice ~ Foraging. **(D)** Across all demonstrator conditions (i.e. the proportion of times any available demonstrator was copied as opposed to making a completely independent decision): Route Choice ~ Foraging. In all cases, each data point represents an individual participant. Regression lines are also shown.

#### 1.1.2. Additional analysis: Success rates of social versus asocial learners in the *Soc/Asoc* condition.

In the main text, we compared the average success rates of participants who tended to learn socially versus those who tended to learn asocially, across all demonstrator conditions. Here, we present the same analysis, but using the *Soc/Asoc* demonstrator condition only – thus comparing the relative successes of participants who tended to copy or ignore a single demonstrator. These results were qualitatively the same as those presented in the main text. In the Same Rewards group, those who tended to learn asocially were significantly more successful than those who favoured social learning in both the Route Choice (Mann-Whitney U test; $U(1) = 265$, $p < 0.001$) and Foraging Tasks (Mann-Whitney U test; $U(1) = 119.5$, $p = 0.012$); whereas in the Different Rewards group, participants who learned largely asocially had similar success rates to those who learned largely socially (Mann-Whitney U tests; Route Choice: $U(1) = 493$, p = 0.295; Foraging: $U(1) = 380$; p = 0.423) (Figure S2). Note that the substantially lower remaining energy in the Route Choice task for the Different Rewards group (Figure S2A) is likely due to the availability of shorter routes in these environments compared to the Same Rewards group and not due to differences in social information use.

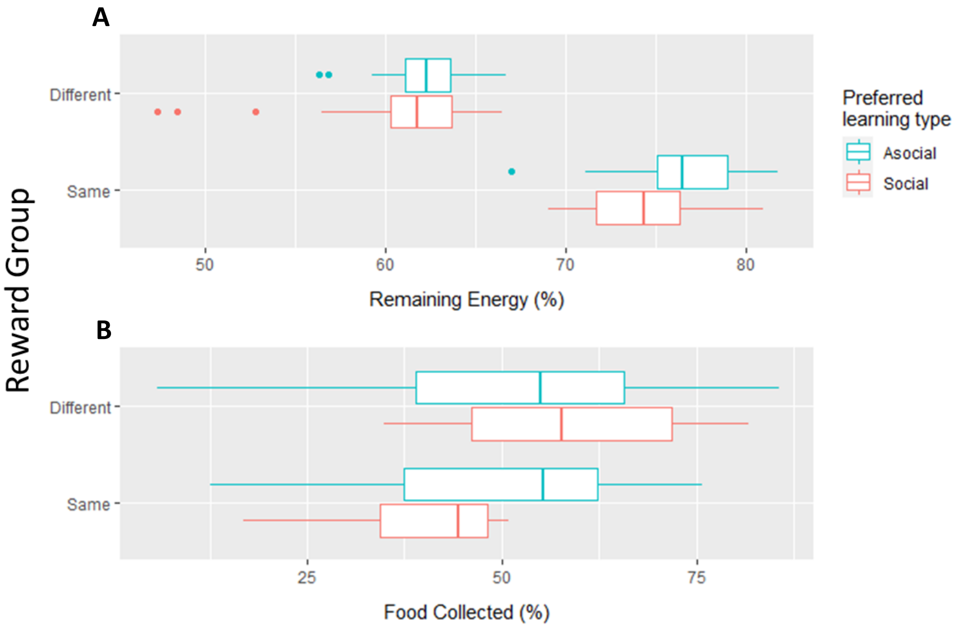

**Figure S2.** The average success rates of participants who favoured social and asocial learning, in the *Soc/Asoc* condition only, in scenarios where demonstrators received different or the same rewards for their choices, across two different tasks. **(A)** In the Route Choice task, success is measured by the average amount of energy remaining when the end point is reached. **(B)** In the Foraging task, success is measured by the average amount of food collected. Boxplots represent the median and interquartile range. Whiskers extend to 1.5x the interquartile range.

#### 1.1.3. Additional analyses of the Foraging task: Did participants learn the specific food type preferences of demonstrators?

In addition to food patch choices, the influence of demonstrator food type choices on participant food collection behaviour was analysed for the Foraging task. For each environment in the Foraging task, food patches were divided into two sections, each containing twenty items of a different food type. Different food types were coloured differently and, in some cases, had different nutritional values. All food types used in the study are given in Table S6. Each demonstrator visited three food patches and 'ate' only one food type from that patch, thus allowing participants to learn preferences for specific food types.

**Table S6.** The food types used in the foraging task, along with their nutritional values.

| **Food item name** | **Appearance** | **Nutritional value** |
|---|---|---|
| *YellowMushroom* | 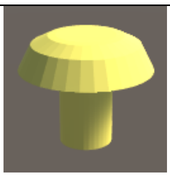 | +5 |
| *PinkMushroom* | 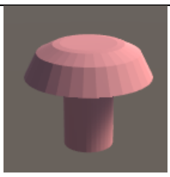 | +3 |
| *BrownMushroom* | 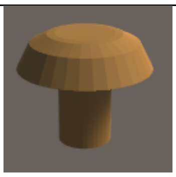 | +1 |
| *GreenMushroom* | 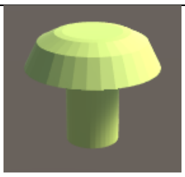 | +1 |
| *WhiteMushroom* | 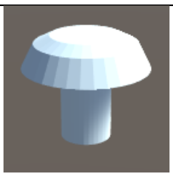 | +1 |
| *DarkMushroom* | 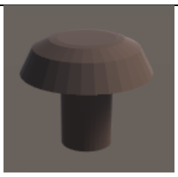 | -5 |

For the Same Rewards group, all eight food patches contained a set of twenty *BrownMushrooms* and a set of twenty *GreenMushrooms* (Table S6). All food items therefore had the same nutritional value of +1. Both demonstrators only ate the *GreenMushrooms* located in their chosen food patches. Thus, if participants were prone to copying the food preferences of certain demonstrators, they should be disproportionately more likely to collect the *GreenMushrooms* over the *BrownMushrooms*. A binomial GLM was run predicting the initial choice made by participants (i.e. the first food item collected during the first demonstrator condition they were subjected to) ($n$ = 57). The reason that initial choices were considered here, rather than the total number of each food type collected by participants, is because AI demonstrators depleted the *GreenMushrooms,* leaving fewer for the player to collect and making it more likely that a GLM would detect a tendency to collect the *BrownMushrooms* more often. According to this model, participants did not show an initial preference towards the demonstrated food type, the *GreenMushrooms* (Table S7).

**Table S7.** Intercept estimates, standard errors, Z values and p-values for GLMs predicting the likelihood that (i) participants in the Same Rewards group would display an initial preference for the demonstrated *GreenMushroom* food type; (ii) participants in the Different Reward group who initially visited the best food patch would display an initial preference for the demonstrated *YellowMushroom* food type; and (iii) participants in the Different Reward group who initially visited the worst food patch would display an initial preference for the demonstrated *WhiteMushroom* food type.

| Reward group | Food patch type | Intercept | Std. Error | Z value | p-value |
|---|---|---:|---:|---:|---:|
| Same Rewards | All | -0.391 | 0.2700 | -1.448 | 0.148 |
| Different Rewards | Best food patch | -0.201 | 0.318 | -0.631 | 0.528 |
| | Worst food patch | 0.154 | 0.556 | 0.277 | 0.782 |

For the Different Rewards group, food patches contained food items of differing nutritional values as follows: Three food patches contained a set of twenty *PinkMushrooms* and a set of twenty *YellowMushrooms* – these were visited by demonstrator A, who ‘ate’ only the *YellowMushrooms* from these patches. Three food patches contained a set of twenty *WhiteMushrooms* and a set of twenty *DarkMushrooms* – these were visited by demonstrator B, who ‘ate’ only the *WhiteMushrooms*. The remaining two (asocial) food patches contained a set of twenty *BrownMushrooms* and a set of twenty *GreenMushrooms*. Thus, demonstrator choices were arranged in such a way that (*i*) copying demonstrator A would result in participants finding the most profitable food patches and (*ii*) copying the food type preference of any demonstrator would result in participants collecting the most profitable food types within a given food patch (and would allow participants to avoid collecting poisonous foods when foraging on the 'worst' food patches). To determine whether participants were more likely to copy the food type preferences of demonstrators in these environments, where rewards were uncertain and potentially maladaptive decisions could be made, two binomial GLMs were run, again using data concerning the initial choice made by each participant. The first model predicted the likelihood that participants who visited the best food patch first ($n$ = 40) also displayed an initial preference towards the food type demonstrated by demonstrator A, i.e. collected the *YellowMushroom* first. The second model predicted the likelihood that participants who visited the worst food patch first ($n$ = 13) also displayed an initial preference towards the food type demonstrated by demonstrator B, i.e. collected the *WhiteMushroom* first. These models revealed that, again, participants were not influenced by the specific food preferences of demonstrators (Table S7). Once they had followed a demonstrator into a food patch, they were equally likely to collect either food type first.

Overall, these analyses suggests that, while certain demonstrators influenced participants' food patch choices, as discussed in the main text, the individual food type preferences of demonstrators were not copied. However, despite this, participants were adaptive in the way they collected food items. When visiting the best food patches, participants collected approximately equal proportions of *PinkMushrooms* and *YellowMushrooms* (Figure S3A). However, when visiting the worst food patches, they disproportionately collected the *WhiteMushrooms* and avoided the poisonous *DarkMushrooms* (Figure S3B). As participants' initial food preferences were unaffected by demonstrator choice, it is likely that participants learned asocially to avoid poisonous foods, probably by sampling them first and learning that this food type lowered the player's health and food score. Participants were therefore adaptive in the way in which they collected food from patches, collecting all food types with a positive nutritional score and ignoring demonstrator food preferences (which would have been maladaptive to copy, since participants would have avoided collecting alternative food types with positive nutritional values in the best food patches), while also avoiding poisonous foods.

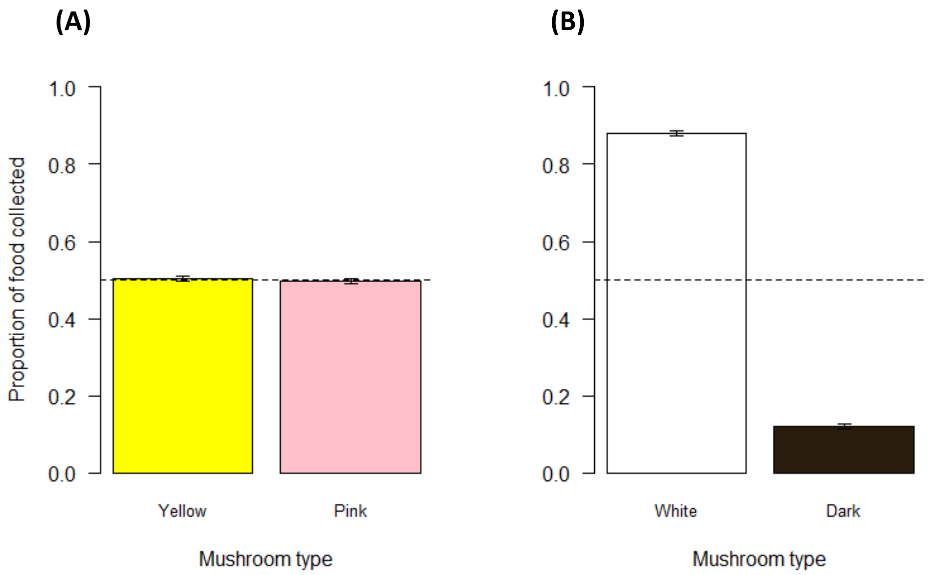

**Figure S3. (A)** The proportion of *YellowMushrooms* (nutritional value: +5) and *PinkMushrooms* (nutritional value: +3) collected by participants when visiting the best food patches. **(B)** The proportion of *WhiteMushrooms* (nutritional value: +1) and *DarkMushrooms* (poisonous; nutritional value: -3) collected when visiting the worst food patches. Proportions are taken from the total number of food items collected across all demonstrator conditions, within the Different Rewards group. Horizontal (dashed) reference line is at 0.5 and indicates no preference for either food type.

#### 1.1.4. Additional analyses concerning participant choices of demonstrator A vs demonstrator B based on their relative success rates

**Table S8.** ANOVA results for GLMs modelling the likelihood of copying demonstrator A over demonstrator B across participants in two different groups – one where rewards were equal regardless of the demonstrator copied, and one where copying demonstrator A over demonstrator B resulted in higher payoffs (as measured by the 'RewardGroup' factor). For each of the three tasks, a null, intercept-only model (representing the likelihood that demonstrator A is copied over B overall, across both reward groups) is compared with a model in which reward group was included as a factor. Significant p-values (< 0.05) are highlighted in bold and indicate significant differences in demonstrator choice across the two reward groups.

| Task | Model | Deviance | Residual Deviance | p-value |
|---|---|---|---|---|
| *Container* | *NULL* | | 603.1 | |
| | *RewardGroup* | 129.9 | 138.0 | **< 0.001** |
| *Route Choice* | *NULL* | | 245.5 | |
| | *RewardGroup* | 26.9 | 218.5 | **< 0.001** |
| *Foraging* | *NULL* | | 433.3 | |
| | *RewardGroup* | 19.6 | 135.0 | **< 0.001** |

**Table S9.** Intercept estimates, standard error, z-values and p-values for binomial GLMs modelling the likelihood that participants copied demonstrator A over demonstrator B in the *Dom/Sub, Three/One, Male/Female* and *Large/Small* demonstrator conditions, and overall across all four demonstrator conditions, for each of the three tasks, in scenarios where payoffs differed depending on which demonstrator was copied. P-values are adjusted for multiple comparisons using false discovery rates. Significant p-values (< 0.05) are highlighted in bold.

| Task | Demonstrator Condition | Intercept Estimate | Std. Error | Z value | p-value |
|---|---|---|---|---|---|
| *Container* | *Dom/Sub* | 0.930 | 0.083 | 11.090 | **<0.001** |
| | *Three/One* | 1.266 | 0.091 | 13.870 | **<0.001** |
| | *Male/Female* | 0.557 | 0.079 | 7.089 | **<0.001** |
| | *Large/Small* | 1.009 | 0.085 | 11.810 | **<0.001** |
| | *All (A/B)* | 0.927 | 0.042 | 22.110 | **<0.001** |

| | | | | | |
|---|---|---|---|---|---|
| *Route Choice* | *Dom/Sub* | 1.263 | 0.214 | 5.900 | **<0.001** |
| | *Three/One* | 1.936 | 0.276 | 7.011 | **<0.001** |
| | *Male/Female* | 0.177 | 0.180 | 0.983 | 0.326 |
| | *Large/Small* | 1.299 | 0.246 | 5.278 | **<0.001** |
| | *All (A/B)* | 1.056 | 0.106 | 10.010 | **<0.001** |
| *Foraging* | *Dom/Sub* | 0.922 | 0.206 | 4.482 | **<0.001** |
| | *Three/One* | 1.266 | 0.197 | 6.421 | **<0.001** |
| | *Male/Female* | 0.786 | 0.188 | 4.173 | **<0.001** |
| | *Large/Small* | 0.936 | 0.212 | 4.414 | **<0.001** |
| | *All (A/B)* | 0.984 | 0.100 | 9.857 | **<0.001** |

#### 1.1.5. Additional analysis – the influence of individual characteristics on their use of social information

A series of binomial GLMs were run to investigate whether participants' individual characteristics influenced their tendency to learn socially over asocially (hypothesis *i* in the *Statistical Analysis,* section 5.3.8, of the main text), their tendency to copy demonstrators with certain characteristics (hypothesis *iv*) and their tendency to copy the most over the least successful demonstrators (hypothesis *v*). The individual characteristics tested were age, gender, aggression score and time spent playing video games. As recommended by Bryant and Smith (2001), the four subcategories of aggression – physical aggression, verbal aggression, hostility and anger – were modelled separately, as these describe different types of aggression that are not necessarily correlated. In addition, the overall aggression score, summed across each of the four categories, was modelled as a measure of general aggression. To avoid issues of collinearity, overall aggression was never included in the same model as any of the four subcategories. For each type of aggression, a participant's score was averaged across all questions in the particular aggression category, and then scaled between 0 and 1.

Time spent playing video games was included in this analysis to assess whether familiarity with video game play influenced participants' general behaviour in VERSE, thus affecting social information use – e.g. if those who play video games often are more exploratory due to higher levels of confidence with video game controls, this may result in a higher tendency to learn asocially. Participants were asked to rate their usual video game usage as either "Never", "Now and again", "A few times a month", "A few times a week" or "Daily". For ease of analysis, these options were then collated into two categories, with the first three options classed as "Rarely" and the remaining two classes as "Often", before insertion into the GLM.

The influence of each individual variable was modelled separately. For each hypothesis to be tested, a binomial GLM was run for each individual variable (age, gender, aggression score, video game play), with said variable as a predictor. Where more than one individual variable was found to have a significant influence on the use of social information, each combination of individual variables were modelled and ANOVA tests and AIC values were used to establish which model provided the best fit to the data.

Overall, there was no clear pattern observed for an influence of the tested individual characteristics on either social information use (Table S10), biases towards certain demonstrator characteristics (Table S11) or tendency to copy more successful demonstrators (Table S12). Most of the models tested were not statistically significant after FDR corrections were performed. However, regarding the influence of age on social learning, it is important to note the relatively narrow age range of the participants (mean: 21; range: 18-31). Different results may have been obtained if a wider age range would have been considered.

Video game play did not appear to have a significant influence on participants' behaviour within VERSE, with one exception: participants who played video games more often were more likely to copy the most successful demonstrators across demonstrator conditions – however this was significant in the Container task only. This could suggest that 'gamers' displayed some different behaviours to 'non-gamers' (e.g. a greater capacity to track the success rates of AIs) during relatively simple, two-option tasks, but not when exposed to more complex, exploratory tasks. All in all, this suggests that, in general, VERSE is well suited to studying human social behaviour in complex, immersive environments irrespective of whether participants are 'gamers' or not, despite VERSE being game-like in its nature.

**Table S10.** Estimates, standard error, z values and p-values obtained from a series of binomial GLMs modelling the influence of each individual characteristic on participants' tendency to use favour social over asocial learning across three tasks, during the *Soc/Asoc* demonstrator condition (where a single demonstrator displayed one option and all other options were undemonstrated) and altogether across all demonstrator conditions. Each individual variable was modelled separately in its own GLM. P-values are adjusted for multiple comparisons using false discovery rates, after which none of the tested models were found to be statistically significant ($p < 0.05$).

| | | **Container** | | | | **Route Choice** | | | | **Foraging** | | | |
|---|---|---|---|---|---|---|---|---|---|---|---|---|---|
| **Individual characteristic** | **Demonstrator condition** | **Estimate** | **Std Error** | **z** | **p** | **Estimate** | **Std Error** | **z** | **p** | **Estimate** | **Std Error** | **z** | **p** |
| Age | *Soc/Asoc* | 0.034 | 0.042 | 0.823 | 0.888 | 0.043 | 0.068 | 0.640 | 0.908 | 0.147 | 0.059 | 2.505 | 0.160 |
| Gender (male) | *Soc/Asoc* | -0.352 | 0.159 | -2.214 | 0.270 | -0.070 | 0.292 | -0.240 | 0.971 | 0.164 | 0.281 | 0.585 | 0.932 |
| Aggression (overall) | *Soc/Asoc* | 0.449 | 0.603 | 0.744 | 0.888 | 0.351 | 1.198 | 0.293 | 0.971 | 0.999 | 1.074 | 0.930 | 0.888 |
| Aggression-physical | *Soc/Asoc* | -0.063 | 0.616 | -0.103 | 0.988 | 0.098 | 1.222 | 0.080 | 0.988 | -0.236 | 1.111 | -0.212 | 0.971 |
| Aggression-verbal | *Soc/Asoc* | 0.079 | 0.397 | 0.198 | 0.971 | 0.005 | 0.751 | 0.007 | 0.995 | 1.958 | 0.697 | 2.809 | 0.100 |
| Aggression-anger | *Soc/Asoc* | 0.933 | 0.447 | 2.085 | 0.296 | 0.269 | 0.863 | 0.312 | 0.971 | -0.313 | 0.820 | -0.382 | 0.971 |
| Aggression-hostility | *Soc/Asoc* | -0.019 | 0.419 | -0.047 | 0.988 | 0.351 | 0.804 | 0.436 | 0.971 | 0.039 | 0.763 | 0.051 | 0.988 |
| Video game play (often) | *Soc/Asoc* | -0.454 | 0.160 | -2.844 | 0.100 | -0.320 | 0.303 | -1.058 | 0.829 | 0.253 | 0.282 | 0.897 | 0.888 |
| Age | *All (soc/asoc)* | - | - | - | - | 0.081 | 0.061 | 1.329 | 0.640 | 0.013 | 0.042 | 0.299 | 0.971 |
| Gender (male) | *All (soc/asoc)* | - | - | - | - | -0.146 | 0.219 | -0.667 | 0.908 | 0.340 | 0.202 | 1.682 | 0.531 |
| Aggression (overall) | *All (soc/asoc)* | - | - | - | - | -1.168 | 0.894 | -1.306 | 0.640 | -0.374 | 0.768 | -0.487 | 0.971 |
| Aggression-physical | *All (soc/asoc)* | - | - | - | - | -0.172 | 0.907 | -0.189 | 0.971 | -1.037 | 0.775 | -1.338 | 0.640 |
| Aggression-verbal | *All (soc/asoc)* | - | - | - | - | -0.418 | 0.553 | -0.757 | 0.888 | 0.748 | 0.497 | 1.507 | 0.640 |
| Aggression-anger | *All (soc/asoc)* | - | - | - | - | -0.904 | 0.625 | -1.447 | 0.640 | -0.174 | 0.564 | -0.308 | 0.971 |
| Aggression-hostility | *All (soc/asoc)* | - | - | - | - | -0.718 | 0.604 | -1.189 | 0.720 | -0.983 | 0.541 | -1.818 | 0.460 |
| Video game play (often) | *All (soc/asoc)* | - | - | - | - | -0.170 | 0.234 | -0.729 | 0.888 | 0.175 | 0.203 | 0.862 | 0.888 |

**Table S11**. Estimates, standard error, z values and p-values obtained from a series of binomial GLMs modelling the influence of each individual characteristic on participants' tendency to copy the dominant demonstrator over the subordinate in the *Dom/Sub* condition, three over one demonstrator in the *Three/One* condition, the male over the female demonstrator in the *Male/Female* condition and the large over the small demonstrator in the *Large/Small* condition, across the three tasks, in conditions where both demonstrator displayed equally profitable behaviours. Each individual variable was modelled separately in its own GLM. P-values are adjusted for multiple comparisons using false discovery rates, after which none of the tested models were found to be statistically significant (p < 0.05).

| | | **Container** | | | | **Route Choice** | | | | **Foraging** | | | |
|---|---|---|---|---|---|---|---|---|---|---|---|---|---|
| **Individual characteristic** | **Demonstrator condition** | **Estimate** | **Std Error** | **z** | **p** | **Estimate** | **Std Error** | **z** | **p** | **Estimate** | **Std Error** | **z** | **p** |
| Age | *Dom/Sub* | 0.024 | 0.042 | 0.587 | 0.945 | -0.089 | 0.132 | -0.673 | 0.945 | 0.021 | 0.091 | 0.234 | 0.945 |
| Gender (male) | *Dom/Sub* | -0.043 | 0.159 | -0.269 | 0.945 | -0.237 | 0.521 | -0.454 | 0.945 | 2.266 | 0.663 | 3.419 | 0.096 |
| Aggression (overall) | *Dom/Sub* | 0.553 | 0.605 | 0.914 | 0.812 | 0.097 | 2.044 | 0.048 | 0.945 | -4.428 | 2.013 | -2.200 | 0.384 |
| Aggression-physical | *Dom/Sub* | -0.243 | 0.617 | -0.394 | 0.945 | 0.593 | 1.824 | 0.325 | 0.945 | -4.761 | 2.036 | -2.339 | 0.365 |
| Aggression-verbal | *Dom/Sub* | -0.083 | 0.398 | -0.210 | 0.945 | 0.639 | 1.348 | 0.474 | 0.945 | -0.064 | 1.360 | -0.047 | 0.995 |
| Aggression-anger | *Dom/Sub* | 0.628 | 0.448 | 1.403 | 0.773 | -0.545 | 1.476 | -0.369 | 0.945 | -3.744 | 1.334 | -2.807 | 0.168 |
| Aggression-hostility | *Dom/Sub* | 0.720 | 0.423 | 1.702 | 0.534 | -0.406 | 1.434 | -0.283 | 0.945 | -1.653 | 1.298 | -1.273 | 0.812 |
| Video game play (often) | *Dom/Sub* | -0.043 | 0.159 | -0.269 | 0.945 | -0.219 | 0.579 | -0.378 | 0.945 | 0.288 | 0.461 | 0.625 | 0.945 |
| Age | *Three/One* | 0.105 | 0.055 | 1.901 | 0.421 | 0.231 | 0.239 | 0.963 | 0.812 | -0.043 | 0.078 | -0.548 | 0.945 |
| Gender (male) | *Three/One* | -0.186 | 0.179 | -1.042 | 0.812 | 0.318 | 0.715 | 0.444 | 0.945 | -0.441 | 0.387 | -1.139 | 0.812 |
| Aggression (overall) | *Three/One* | 0.697 | 0.694 | 1.004 | 0.812 | -2.182 | 2.649 | -0.824 | 0.856 | 3.170 | 1.604 | 1.976 | 0.400 |
| Aggression-physical | *Three/One* | -1.034 | 0.668 | -1.549 | 0.645 | -2.552 | 2.670 | -0.956 | 0.812 | -0.109 | 1.512 | -0.072 | 0.995 |
| Aggression-verbal | *Three/One* | 0.460 | 0.458 | 1.006 | 0.812 | -0.713 | 1.544 | -0.462 | 0.945 | 1.564 | 0.961 | 1.628 | 0.582 |
| Aggression-anger | *Three/One* | 0.312 | 0.511 | 0.612 | 0.945 | -2.256 | 1.694 | -1.331 | 0.793 | 3.458 | 1.277 | 2.708 | 0.168 |
| Aggression-hostility | *Three/One* | 1.091 | 0.495 | 2.204 | 0.384 | 0.473 | 1.746 | 0.271 | 0.945 | 1.265 | 1.089 | 1.161 | 0.812 |
| Video game play (often) | *Three/One* | 0.172 | 0.183 | 0.940 | 0.812 | 0.337 | 0.840 | 0.401 | 0.945 | 0.096 | 0.422 | 0.228 | 0.945 |

| | | | | | | | | | | | | | |
|---|---|---|---|---|---|---|---|---|---|---|---|---|---|
| Age | *Male/Female* | 0.002 | 0.041 | 0.041 | 0.995 | 0.200 | 0.180 | 1.113 | 0.811 | 0.050 | 0.103 | 0.489 | 0.945 |
| Gender (male) | *Male/Female* | 0.000 | 0.158 | 0.000 | 1.000 | -0.309 | 0.511 | -0.604 | 0.945 | -0.388 | 0.402 | -0.965 | 0.812 |
| Aggression (overall) | *Male/Female* | 0.874 | 0.605 | 1.445 | 0.748 | -2.077 | 1.947 | -1.066 | 0.812 | -1.352 | 1.573 | -0.859 | 0.832 |
| Aggression-physical | *Male/Female* | 0.127 | 0.616 | 0.205 | 0.945 | -0.027 | 1.810 | -0.015 | 0.998 | 0.095 | 1.575 | 0.060 | 0.995 |
| Aggression-verbal | *Male/Female* | -0.263 | 0.397 | -0.661 | 0.945 | -1.595 | 1.209 | -1.320 | 0.793 | -0.285 | 1.027 | -0.277 | 0.945 |
| Aggression-anger | *Male/Female* | 0.878 | 0.447 | 1.963 | 0.400 | -1.375 | 1.466 | -0.938 | 0.812 | -0.286 | 1.155 | -0.248 | 0.945 |
| Aggression-hostility | *Male/Female* | 1.151 | 0.425 | 2.708 | 0.168 | -0.810 | 1.337 | -0.606 | 0.945 | -2.512 | 1.236 | -2.033 | 0.400 |
| Video game play (often) | *Male/Female* | -0.050 | 0.158 | -0.317 | 0.995 | -0.474 | 0.527 | -0.899 | 0.812 | 0.013 | 0.419 | 0.032 | 0.995 |
| Age | *Large/Small* | 0.002 | 0.041 | 0.043 | 0.995 | 0.081 | 0.102 | 0.798 | 0.868 | -0.280 | 0.140 | -2.001 | 0.400 |
| Gender (male) | *Large/Small* | -0.270 | 0.159 | -1.700 | 0.534 | -0.509 | 0.410 | -1.243 | 0.812 | 0.507 | 0.442 | 1.147 | 0.812 |
| Aggression (overall) | *Large/Small* | 0.792 | 0.604 | 1.311 | 0.793 | -0.725 | 1.701 | -0.426 | 0.945 | -1.432 | 1.575 | -0.909 | 0.812 |
| Aggression-physical | *Large/Small* | 0.344 | 0.617 | 0.558 | 0.945 | -1.329 | 2.196 | -0.605 | 0.945 | -1.612 | 1.805 | -0.893 | 0.812 |
| Aggression-verbal | *Large/Small* | 0.423 | 0.398 | 1.065 | 0.812 | 0.548 | 1.083 | 0.506 | 0.945 | -0.061 | 1.037 | -0.059 | 0.995 |
| Aggression-anger | *Large/Small* | 0.914 | 0.447 | 2.047 | 0.400 | -0.500 | 1.249 | -0.400 | 0.945 | -0.087 | 1.133 | -0.077 | 0.995 |
| Aggression-hostility | *Large/Small* | 0.086 | 0.419 | 0.205 | 0.945 | -1.088 | 1.125 | -0.967 | 0.812 | -2.015 | 1.118 | -1.802 | 0.494 |
| Video game play (often) | *Large/Small* | -0.019 | 0.158 | -0.117 | 0.995 | -0.266 | 0.427 | -0.624 | 0.945 | -0.160 | 0.433 | -0.370 | 0.945 |

**Table S12.** Estimates, standard error, z values and p-values for the interaction between the individual characteristics of participants and the 'reward group' they were assigned to, produced from binomial GLMs modelling the likelihood that participants copied demonstrator A over demonstrator B across all demonstrator conditions. Reward group is a binomial factor describing whether demonstrators received the same or different rewards for their actions (reference group = same). The interaction term therefore describes how much more likely participants with particular characteristics were to copy more successful demonstrators, relative to any innate preferences for the individual characteristics of those demonstrators. P-values are adjusted for multiple comparisons using false discovery rates. Statistically significant estimates ($p < 0.05$) are highlighted in bold and significant models are additionally highlighted in green.

| | **Container** | | | | **Route Choice** | | | | **Foraging** | | | |
|---|---|---|---|---|---|---|---|---|---|---|---|---|
| **Individual characteristic** | **Estimate** | **Std Error** | **z** | **p** | **Estimate** | **Std Error** | **z** | **p** | **Estimate** | **Std Error** | **z** | **p** |
| Age | -0.095 | 0.035 | -2.688 | 0.084 | -0.053 | 0.103 | -0.512 | 0.745 | -0.071 | 0.077 | -0.911 | 0.579 |
| Gender (male) | 0.254 | 0.120 | 2.114 | 0.202 | 0.445 | 0.344 | 1.293 | 0.465 | -0.667 | 0.295 | -2.260 | 0.192 |
| Aggression (overall) | -0.086 | 0.451 | -0.192 | 0.885 | -0.140 | 1.293 | -0.108 | 0.914 | 1.781 | 1.119 | 1.592 | 0.331 |
| Aggression-physical | 0.507 | 0.442 | 1.146 | 0.465 | -0.658 | 1.334 | -0.493 | 0.746 | 1.613 | 1.048 | 1.540 | 0.331 |
| Aggression-verbal | 0.479 | 0.307 | 1.562 | 0.331 | -0.985 | 0.845 | -1.166 | 0.465 | 0.188 | 0.758 | 0.248 | 0.877 |
| Aggression-anger | -0.254 | 0.327 | -0.778 | 0.656 | 0.403 | 0.906 | 0.445 | 0.750 | 0.985 | 0.793 | 1.242 | 0.465 |
| Aggression-hostility | -0.664 | 0.327 | -2.030 | 0.202 | 0.571 | 0.916 | 0.624 | 0.711 | 1.538 | 0.800 | 1.923 | 0.216 |
| Video game play (often) | 0.501 | 0.127 | 3.934 | **0.002** | 0.206 | 0.347 | 0.594 | 0.711 | 0.310 | 0.305 | 1.015 | 0.531 |
